# mTOR-neuropeptide Y signaling sensitizes nociceptors to drive neuropathic pain

**DOI:** 10.1101/2021.10.29.466458

**Authors:** Lunhao Chen, Yaling Hu, Siyuan Wang, Kelei Cao, Weihao Mai, Weilin Sha, Huan Ma, Yong-Jing Gao, Shumin Duan, Yue Wang, Zhihua Gao

**Author notes:** Correspondence (Z.G.) and (Y. W.). These authors contributed equally to this work (Lunhao Chen, Yaling Hu, Siyuan Wang).

## Abstract

Neuropathic pain is a refractory condition that involves de novo protein synthesis in the nociceptive pathway. The mechanistic target of rapamycin (mTOR) is a master regulator of protein translation; however, mechanisms underlying its role in neuropathic pain remain elusive. Using spared nerve injury-induced neuropathic pain model, we found that mTOR is preferentially activated in large-diameter dorsal root ganglion (DRG) neurons and spinal microglia. However, selective ablation of mTOR in DRG neurons, rather than microglia, alleviated neuropathic pain. We show that injury-induced mTOR activation promoted transcriptional induction of NPY likely via signal transducer and activator of transcription 3 (STAT3) phosphorylation. NPY further acted primarily on Y2 receptors (Y2R) to enhance nociceptor excitability. Peripheral replenishment of NPY reversed pain alleviation upon mTOR removal, whereas Y2R antagonists prevented pain restoration. Our findings reveal an unexpected link between mTOR and NPY in promoting nociceptor sensitization and neuropathic pain, through NPY/Y2R signaling-mediated intra-ganglionic transmission.

## INTRODUCTION

Chronic pain, the leading cause of long-term human disability, poses a heavy health burden to the society. Nerve injury-induced neuropathic pain accounts for approximately one fifth of the chronic pain population (van Hecke et al., 2014). It is characterized by persistent hyperalgesia, allodynia and spontaneous pain. Long-lasting sensitization of the nociceptive pathway, leading to a reduced pain threshold, has been considered a major mechanism mediating the persistent hypersensitivity in neuropathic pain (Costigan et al., 2009).

Accumulating evidence has shown that nerve injury-induced *de novo* gene expression contributes to the maladaptive responses in both the peripheral and central nociceptive circuits, thereby promoting nociceptive sensitization and pain hypersensitivity (Costigan et al., 2009; Melemedjian and Khoutorsky, 2015). Elevation of G protein-coupled receptors (GPCRs), such as GPR151, coupled with ion channels in the injured dorsal root ganglion (DRG), has been shown to facilitate the generation of ectopic action potential in nociceptive neurons and promotes pain (Geppetti et al., 2015; Xia et al., 2021).

Other than ion channels and GPCRs, prominent induction of neuropeptides, including neuropeptide Y (NPY), galanin (Gal), neurotensin (NTS) and cholecystokinin (CCK), have also been observed in DRG neurons after nerve injury (Reinhold et al., 2015; Wu et al., 2016; Xiao et al., 2002). The 36-amino acid peptide, NPY, is one of the most robustly upregulated neuropeptides in DRG neurons after nerve injury (Wakisaka et al., 1991). However, mechanisms underlying its induction remain unknown. Moreover, conditional knockdown of spinal cord NPY increased tactile and thermal hypersensitivity primarily through Y1 receptor (Y1R) in nerve injury-induced neuropathic pain models (Solway et al., 2011; Nelson and Taylor, 2021), whereas subcutaneous injection of NPY or Y2 receptor (Y2R) agonist exacerbated pain after nerve injury, suggesting a biphasic role of NPY in neuropathic pain at different sites (Sapunar et al., 2011; Tracey et al., 1995; Arcourt et al., 2017). It remains to be elucidated how NPY was induced after injury and whether NPY plays opposing roles through different receptors in the nociceptive pathway.

The mechanistic target of rapamycin (mTOR), a master regulator of protein translation, plays a pivotal role in regulating cell growth and metabolism. Deregulation of mTOR signaling has been linked to various human diseases, including cancer, obesity and neurodegeneration (Saxton and Sabatini, 2017; Carlin et al., 2018; Laplante and Sabatini, 2013). Activation of mTOR has been observed in the DRG and spinal cord in neuropathic pain models, as well as morphine-induced chronic pain (Abe et al., 2010; Zhang et al., 2013; Xu et al., 2014; Melemedjian et al., 2011; Price and Géranton, 2009). Pharmacologic blockade of mTOR activity has been demonstrated to reduce pain (Geranton et al., 2009; Asante et al., 2010; Obara et al., 2011; Tateda et al., 2017; Norsted Gregory et al., 2010; Xu et al., 2011). However, several studies found that inhibiting mTOR complex 1 (mTORC1) resulted in unexpected mechanical allodynia, through an insulin receptor substrate-1 (IRS-1)-dependent negative feedback activation of extracellular signal-regulated kinase (ERK) in primary sensory neurons (Melemedjian et al., 2013; Melemedjian and Khoutorsky, 2015), leaving the role of mTOR and underlying mechanisms in pain regulation to be further clarified.

Combining genetic manipulation, transcriptomic profiling with electrophysiological recording, we uncover a previously unrecognized link between nerve injury-triggered mTOR activation and NPY induction in DRG neurons. We further demonstrate that mTOR-mediated NPY production enhances nociceptor excitability and promotes pain hypersensitivity through Y2R in DRG. Although mTOR-related signaling has been extensively studied, we present the first evidence for mTOR-regulated NPY signaling in driving neuropathic pain development.

## RESULTS

### Nerve injury induces mTOR activation in subsets of DRG neurons and spinal cord microglia

To examine the status of mTOR activation after nerve injury, we carried out western blot analysis of L4 and L5 DRG and spinal dorsal horn (SDH) tissues from mice at different time points after the spared nerve injury (SNI) surgery (**Figure 1A**). The activity of mTOR was assessed by the levels of phosphorylated S6 protein (p-S6), a key and downstream effector of mTOR. As shown in **Figure 1B and C**, p-S6 was substantially upregulated in the ipsilateral DRG one day after the nerve injury and lasted for at least 7 days (*p*<0.05). These data are consistent with elevated mTOR activity in DRGs after peripheral nerve injury (Abe et al., 2010).

**Figure 1.**
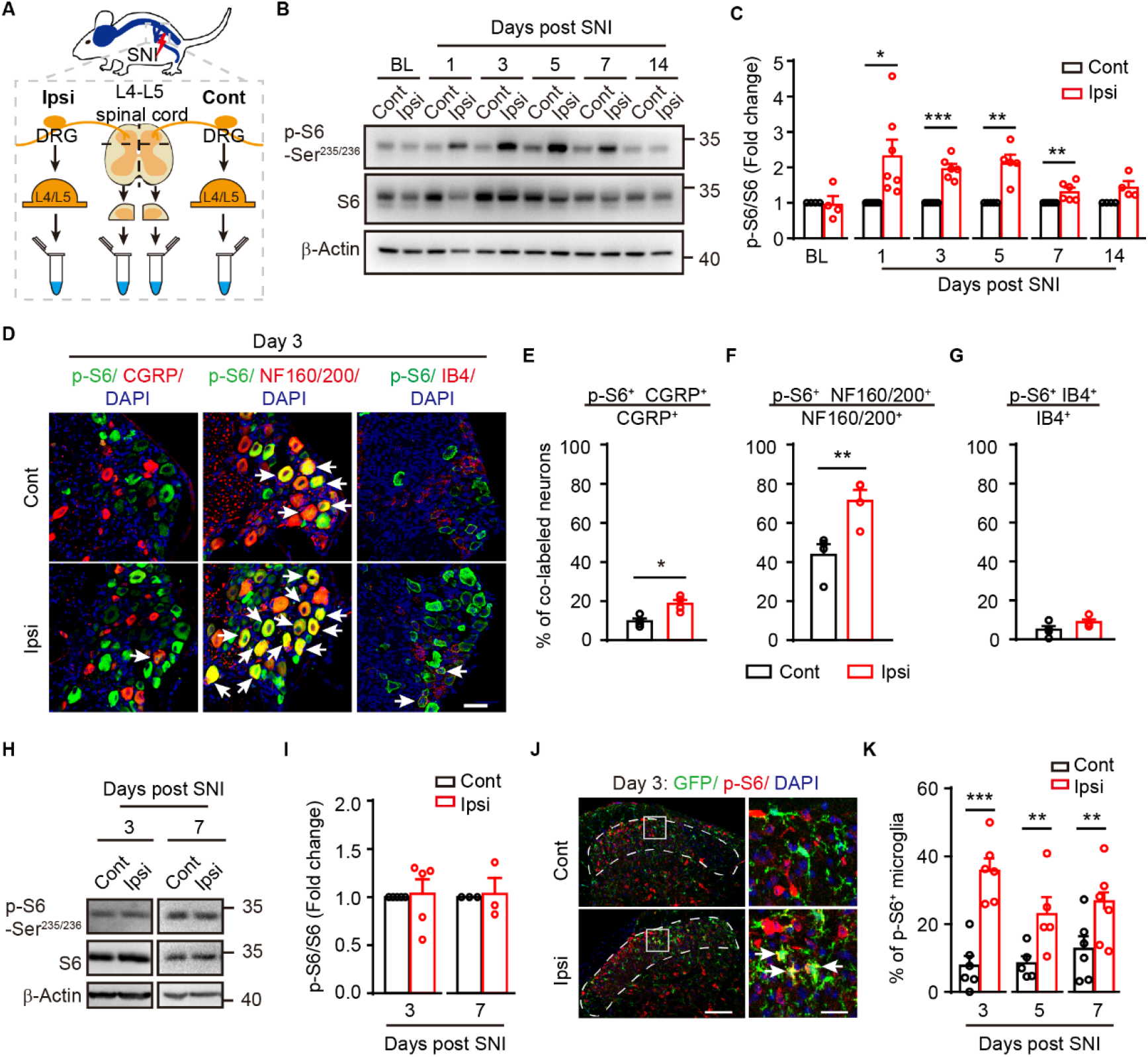
Activation of the mTOR in subsets of DRG neurons and SDH microglia after spared nerve injury (SNI). (**A**) A schematic diagram depicting the isolation of DRGs and SDH. (**B**) Representative blots indicating the upregulated phosphor-S6 (p-S6) levels in the ipsilateral DRGs after SNI. (**C**) Quantification of p-S6/S6 in ipsilateral DRG at indicated time points after SNI (*n* = 4-7 mice per time point). (**D**) Co-immunostaining p-S6 with CGRP, NF160/200 or IB4 in DRGs after SNI (arrows indicating co-labeled neurons). Scale bar, 50 μm. (**E-G**) Quantification of p-S6^+^ neurons in different subpopulations of DRG neurons: CGRP **(E**), NF160/200 (**F**), and IB4 (**G**) (*n* = 4 mice). (**H**) Representative blots of p-S6 and S6 levels in SDH (L4-L5) at day 3 and day 7 after SNI. (**I**) Quantification of p-S6/S6 in ipsilateral and contralateral SDH (*n* = 5 and 3 for day 3 and day 7 post SNI respectively). (**J**) Representative images of p-S6^+^ microglia (arrows) in superficial contralateral and ipsilateral SDH (dotted lines) at indicated time points after SNI. Boxes show regions of higher magnification in the SDH. Scale bar, 100 μm for low magnification images and 20 μm for high magnification images. (**K**) Quantification of p-S6^+^ microglia in superficial SDH (*n* = 5-6 mice per time point). Values are means ± SEM. * *p*<0.05, ** *p*<0.01, and *** *p*<0.001, paired student t-tests. BL, baseline; Ipsi, ipsilateral; Cont, contralateral; DRG: dorsal root ganglion; SDH, spinal dorsal horn. **Figure 1-source data 1.** Raw data of quantification of p-S6/S6 blots, p-S6^+^ neurons, p- S6^+^ microglia in DRG and SDH. **Figure 1-source data 2.** Original pictures of the western blots presented in (B) and (H). **Figure 1-figure supplement 1.** Characterizing of p-S6 staining with different markers in DRG or SDH after SNI.

To further determine the identity of cells with mTOR activation, we performed immunofluorescence analysis using anti-p-S6 antibody along with different markers. Size frequency analysis showed that expression of p-S6 in DRG neurons was mainly in medium and large sizes at contralateral and ipsilateral DRGs after SNI (**Figure 1-figure supplement 1A**). In the contralateral DRG, positive p-S6 labeling, reflecting basal mTOR activity, was observed in a small subset of CGRP^+^ peptidergic neurons (9.7%) but a large fraction of NF160/200^+^ neurons, reminiscent of large-sized A-fiber mechanoreceptors (43.7%). In the ipsilateral DRG, a substantial increase of p-S6^+^ cells in NF160/200^+^ large-sized mechanoreceptors (from 43.7% to 71.2%, *p*<0.01) and CGRP^+^ peptidergic neurons (from 9.7% to 18.7%, *p*<0.05) was observed 3 days after SNI (**Figure 1D-F**). Notably, no elevation of mTOR activity was observed in IB4^+^ non- peptidergic small neurons (*p*>0.05, **Figure 1G**).

By contrast, western blot analysis of p-S6 from the SDH tissue extracts detected no differences between the contralateral and ipsilateral spinal cords following SNI (*p*>0.05, **Figure 1H and I**). Given that western blot analysis detects the gross mTOR activity in the SDH, which may mask changes in sparsely distributed cells in the spinal cord, we carried out dual-labeling of p-S6 with different cellular markers, including NeuN (neurons), GFAP (astrocytes) and Iba1 (microglia). No changes were observed in p-S6^+^ neurons or astrocytes between the contralateral and ipsilateral SDH within 1 week following the injury (**Figure 1-figure supplement 1B-D**). However, the number of p- S6^+^ microglia (GFP^+^) in the superficial layers of ipsilateral SDH was robustly increased from day 3 to 7 post SNI in *Cx3cr1^EGFP/+^* mice (*p<0.05*, **Figure 1J and K**). Together, our results demonstrate that peripheral nerve injury induces mTOR activation mainly in large-sized DRG mechanoreceptors and SDH microglia.

### Blocking mTOR activity delays pain development

To further determine the contribution of mTOR signaling in neuropathic pain, we administered rapamycin, an mTORC1 inhibitor, by intraperitoneal injection to systematically blocking the mTORC1 activity. Meanwhile, BrdU was injected into the mice to label proliferating microglia (**Figure 2A**). Daily intraperitoneal administration of rapamycin from one day before to 7 days after the SNI significantly inhibited mTOR activity in both DRG neurons and SDH microglia (**Figure 2-figure supplement 1**). Using von Frey tests, we found that systemic rapamycin administration delayed the appearance of mechanical allodynia for 5 days after the nerve injury (*p*<0.05, **Figure 2B**). Rapamycin treatment also reduced the total number of microglia (Vehicle: 757.7 ± 15.4 per mm^2^, Rapamycin: 463.1 ± 20.7 per mm^2^, *p<0.001*) (**Figure 2C and D**) and the percentage of proliferative microglia (BrdU^+^ Iba1^+^) (Vehicle: 86.9% ± 1.1%, Rapamycin: 73.1% ± 2.1%) in the superficial layers of ipsilateral SDH at day 7 after SNI (**Figure 2C and E**). These data demonstrated that blocking mTOR signaling delayed pain and suppressed nerve damage-induced microgliosis.

**Figure 2.**
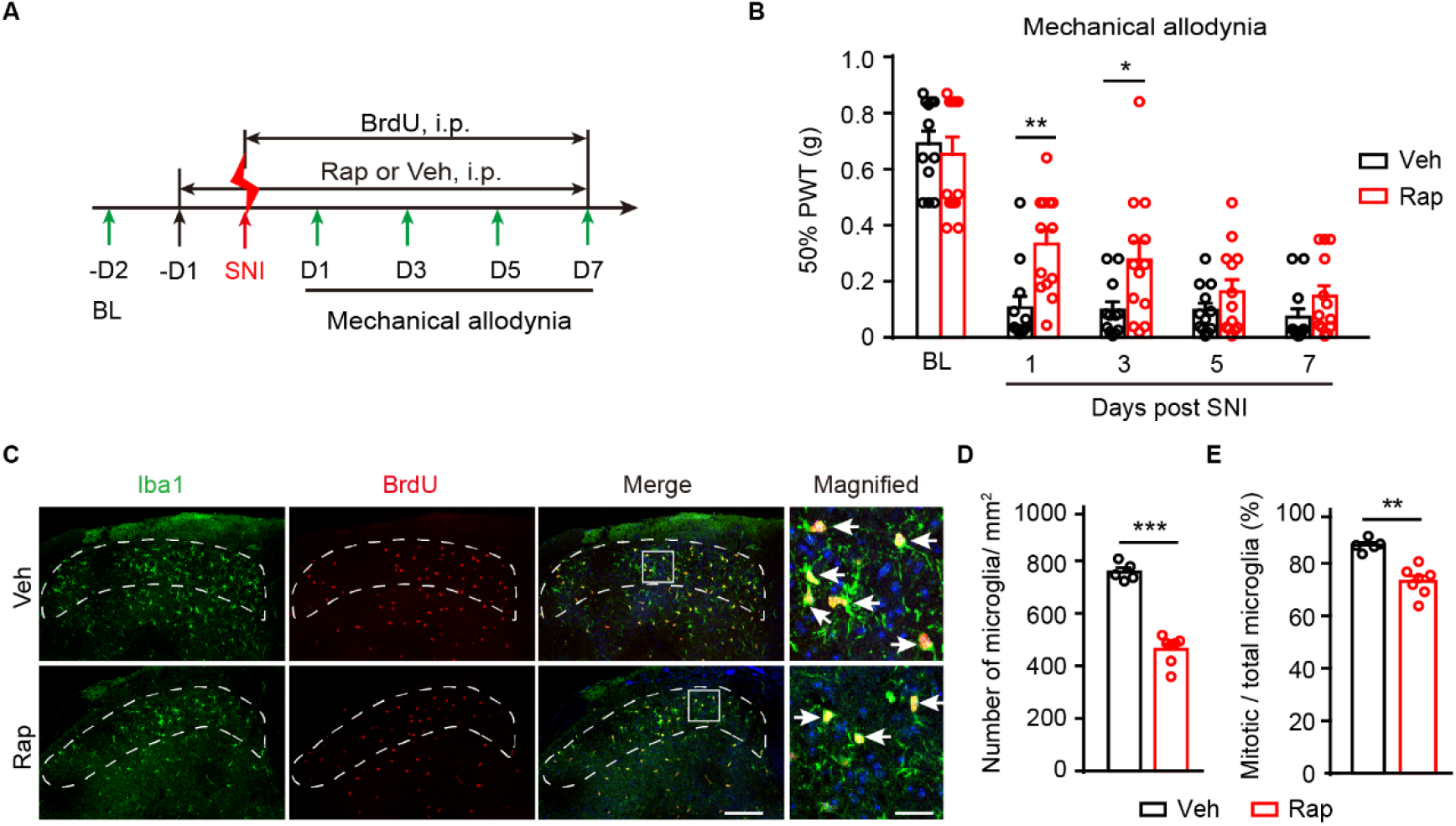
Rapamycin treatments inhibit mTOR activation and attenuates mechanical allodynia after SNI. (**A**) Experimental schedule for rapamycin or vehicle administration through intraperitoneal (i.p.) injection. (**B**) Measurements of mechanical allodynia with daily i.p. injection of rapamycin or vehicle after SNI (*n* = 12-13 per group). (**C**) Representative images of Iba1 and BrdU immunolabeling in ipsilateral superficial SDH (dotted regions) at day 7 after SNI. Boxes show regions of higher magnification in SDH, while arrows indicate Iba1^+^ BrdU^+^ mitotic microglia. Scale bars, 100 μm for low magnification images and 20 μm for high magnification images. (**D-E**) Quantitative analysis of microglia per square millimeter (**D**) and the percentage of mitotic microglia in total microglia (**E**) in ipsilateral SDH at day 7 after SNI (*n* = 5- 7 mice per group). Values are means ± SEM. * *p*<0.05, ** *p*<0.01, *** *p*<0.001, two-way ANOVA followed by Bonferroni’s *post hoc* tests among group (**B**), or unpaired student t-tests (**D, E**). Rap, rapamycin; Veh, vehicle; BL, baseline; D, day; SDH, spinal dorsal horn; PWT, paw withdraw threshold. **Figure 2-source data 1.** Raw data of measurements of mechanical allodynia, number of microglia per square millimeter, and the percentage of mitotic microglia in total microglia. **Figure 2-figure supplement 1.** Administration of rapamycin suppresses mTOR activation.

### Selective ablation of mTOR in DRG neurons but not in microglia alleviates neuropathic pain

To further discern the contributions of neuronal or microglial mTOR in neuropathic pain, we crossed specific Cre mouse lines (*Adv^cre^* or *Cx3cr1^creER^*) with *Mtor^fl/fl^* mice to selectively delete *Mtor* gene in primary sensory neurons or microglia, respectively. We observed complete elimination of p-S6 in DRG neurons and unchanged p-S6 levels in SDH in *Adv^cre^::Mtor^fl/fl^* (*Mtor-cKO^Adv^*) mice 7 days after SNI (**Figure 3A and B**), demonstrating the selective removal of *Mtor* in primary sensory neurons. Examination of sensory perception and motor activities found no significant differences between the control and *Mtor-cKO^Adv^* mice at basal states (**Figure 3-figure supplement 1**). However, *Mtor-cKO^Adv^* mice exhibited delayed development of mechanical allodynia (**Figure 3C**) and cold allodynia (**Figure 3E**) than the controls after SNI, as well as alleviated heat hyperalgesia (**Figure 3D**). Moreover, *Mtor-cKO^Adv^* mice had lower difference scores in response to mechanical stimulation than the *Mtor^fl/fl^* mice in a two- chamber CPA assay that assesses the aversive responses to pain, suggesting that mTOR deletion in DRG neurons alleviated aversive responses to noxious stimuli (**Figure 3F**).

**Figure 3.**
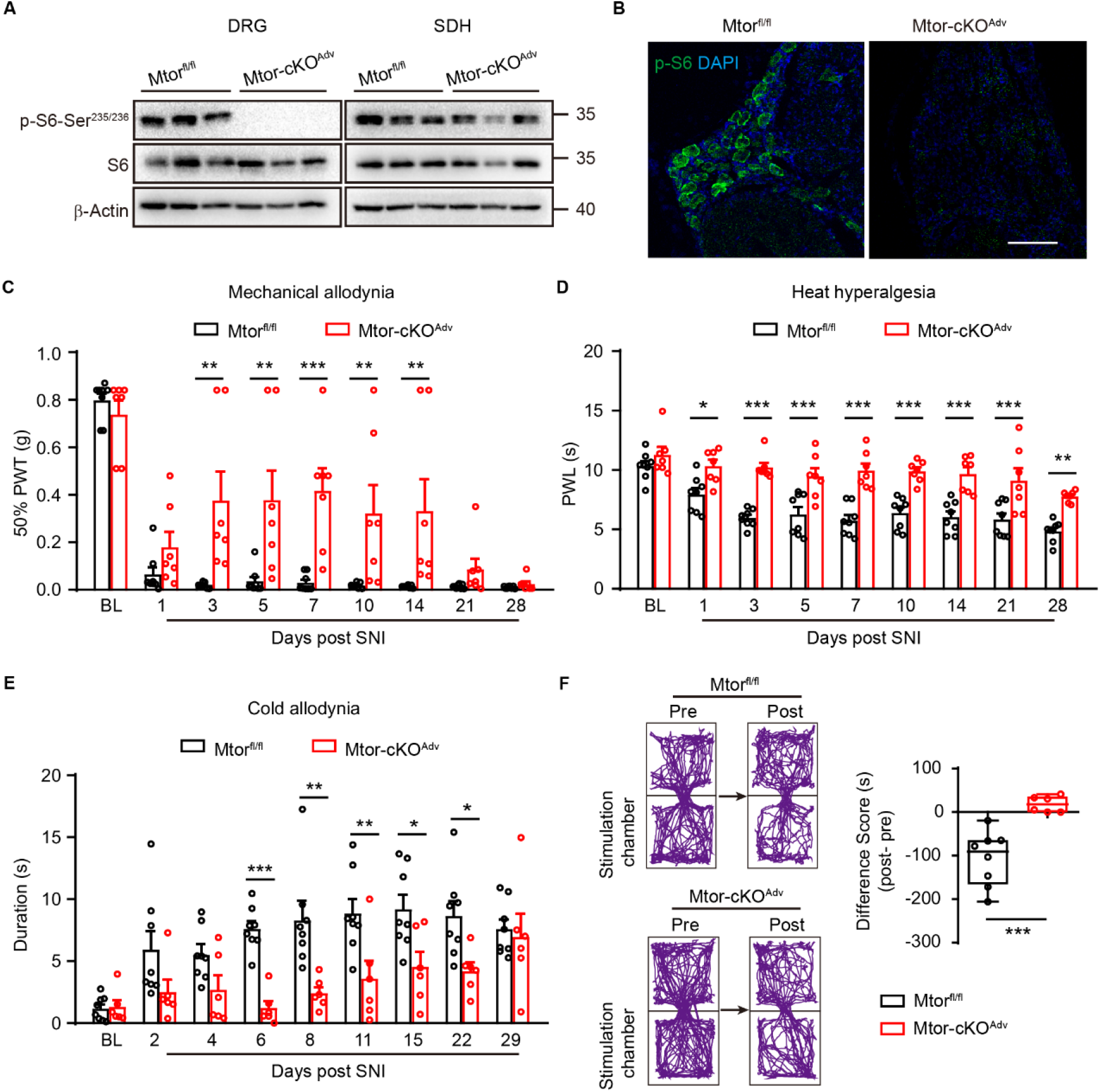
Ablation of *Mtor* in DRG neurons alleviates neuropathic pain. (**A**) Representative blots of p-S6 and S6 in ipsilateral DRG and SDH from *Mtor^fl/fl^* and *Mtor-cKO^Adv^* mice at day 7 after SNI. (**B**) Representative images of p-S6 in ipsilateral DRG at day 7 after SNI, indicating the ablation of mTOR in *Mtor-cKO^Adv^* mice rather than *Mtor^fl/fl^* mice after SNI. Scale bar, 100 μm. (**C-E**) Measurements of mechanical allodynia (**C**), heat hyperalgesia (**D**), and cold allodynia (**E**) in *Mtor^fl/fl^* and *Mtor-cKO^Adv^* mice before and after SNI (*n* = 6-8 mice per group). (**F**) Track plots of animal movements at pre- and post-conditioning phase with a two-chamber conditioned place aversion (CPA) test (*n* = 6-8 mice per group) in *Mtor^fl/fl^* and *Mtor-cKO^Adv^* mice at day 15 after SNI. Difference scores = post-conditioning time (post) - pre-conditioning (pre) time spent in the stimulation chamber. Values are means ± SEM. * *p*<0.05, ** *p*<0.01, *** *p*<0.001, two-way ANOVA followed by Bonferroni’s *post hoc* tests among groups (**C-E**), or unpaired student t-tests (**F**). BL, baseline, PWT, paw withdraw threshold; PWL, paw withdraw latency. **Figure 3-source data 1.** Raw data of measurements of mechanical allodynia, heat hyperalgesia, cold allodynia, and CPA tests. **Figure 3-source data 2.** Original pictures of the western blots presented in (A). **Figure 3-figure supplement 1.** Sensory functions and motor activities are comparable in *Mtor^fl/fl^* and *Mtor-cKO^Adv^* mice at basal state.

To further examine whether microglial mTOR activation also contributes to neuropathic pain, we selectively deleted *Mtor* in microglia by injecting tamoxifen into the *Cx3cr1^creER/+^::Mtor^fl/fl^* mice (*Mtor-cKO^MG^* mice) 4-6 weeks before the SNI surgery (**Figure 4A** **and Figure 4-figure supplement 1A**) (Gu et al., 2016). Cre-mediated recombination of *Mtor* gene in the central nervous system (brain and spinal cord) was detected by PCR analysis (**Figure 4-figure supplement 1B**) and ablation of mTOR in microglia was verified by immunofluorescence analysis (**Figure 4B**). At day 7 post SNI, we observed a reduction in the number of microglia (**Figure 4C and D**) and the percentage of mitotic microglia (BrdU^+^ Iba1^+^) (**Figure 4E and F**) in the superficial layers of ipsilateral SDH in *Mtor-cKO^MG^* mice. However, we were unable to observe significant differences in mechanical allodynia (**Figure 4G**) or heat hyperalgesia (**Figure 4H**) between the *Mtor-cKO^MG^* and control mice after SNI (from day 1 to day 7), suggesting that neuropathic pain is spared in the absence of microglial mTOR signaling.

**Figure 4.**
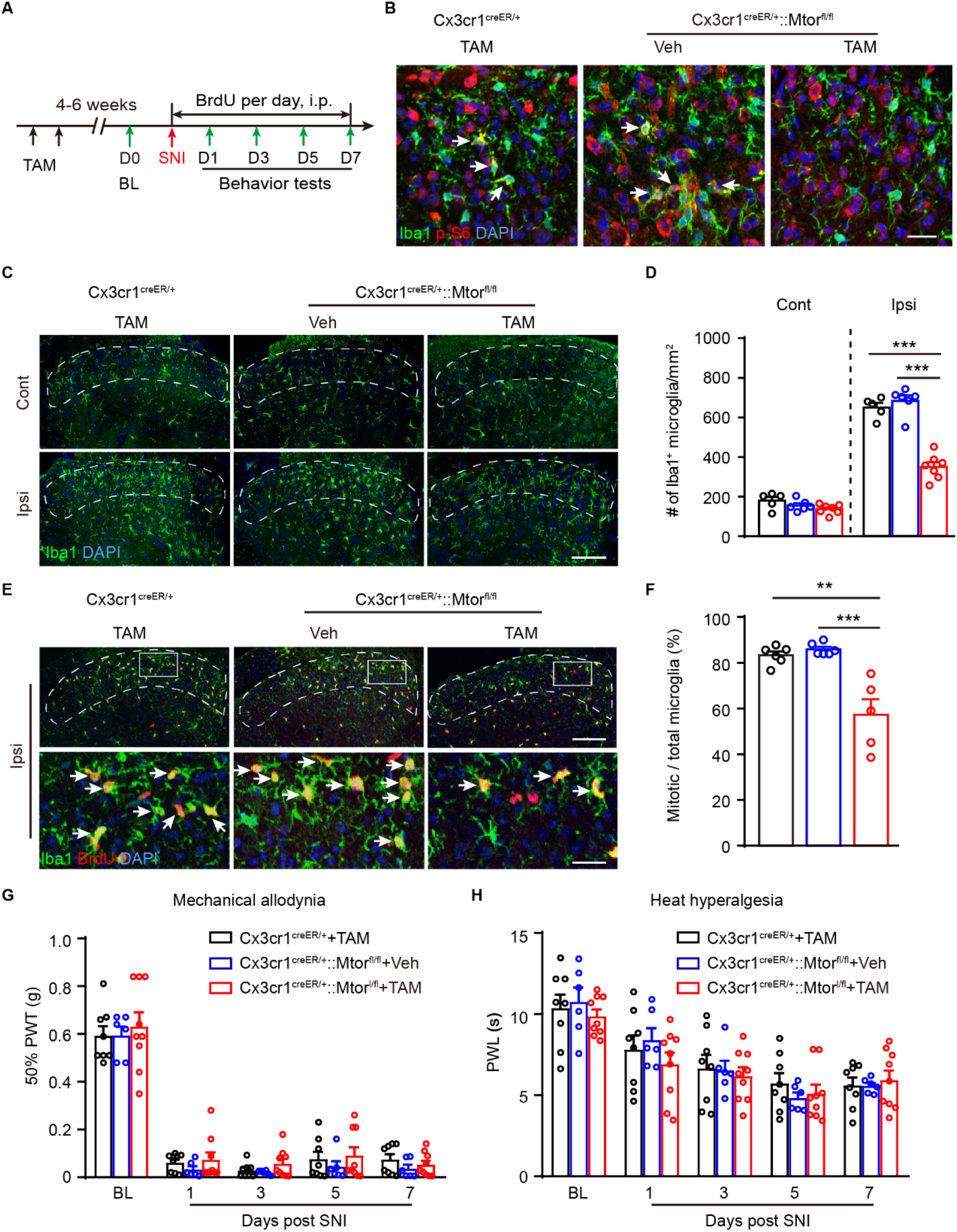
Ablation of *Mtor* in microglia reduces microgliosis, but does not affect neuropathic pain. (**A**) Experimental schedule showing the selected *Mtor* deletion in microglia and pain tests. (**B**) Representative images showing immunofluorescence labeling of Iba1 and p-S6 in ipsilateral SDH at day 7 post SNI in *Cx3cr1^CreER/+^::Mtor^fl/fl^* or control mice (*Cx3cr1^CreER/+^* mice with TAM and *Cx3cr1^CreER/+^::Mtor^fl/fl^* mice with Veh). Arrows indicating Iba1^+^ p-S6^+^ microglia. Scale bar, 20 μm. (**C**) Representative images of bilateral SDH microglia (Iba1^+^) in *Cx3cr1^CreER/+^::Mtor^fl/fl^* mice with TAM or in control mice at day 7 after SNI. Scale bar, 100 μm. (**D**) Quantification of microglia in ipsilateral and contralateral in *Cx3cr1^CreER/+^::Mtor^fl/fl^* and control mice at day 7 post SNI (*n* = 5-7 per group). (**E**) Representative images of ipsilateral SDH showing co-localization of Iba1 and BrdU (arrows) at day 7 after SNI. Boxes show regions of higher magnification in the SDH. Scale bar, 100 μm for low magnification images and 20 μm for high magnification images. (**F**) Quantitation of mitotic microglia (Iba1^+^ BrdU^+^) in SDH in *Cx3cr1^CreER/+^::Mtor^fl/fl^* and control mice at day 7 after SNI (*n* = 5-7 mice per group). (**G-H**) Measurements of mechanical allodynia (**G**) and heat hyperalgesia (**H**) in *Cx3cr1^CreER/+^::Mtor^fl/fl^* and control mice before and after SNI (*n* = 6-9 mice per group). Values are means ± SEM. ** *p*<0.01, and *** *p*<0.001, one-way AVOVA (**F**) or two-way ANOVA followed by Bonferroni’s *post hoc* tests among groups (**D, G, H**). TAM, tamoxifen; Veh, vehicle; Cont, contralateral; Ipsi, ipsilateral; PWT, paw withdraw threshold; PWL, paw withdraw latency; D, day. **Figure 4-source data 1.** Raw data of quantification of microglia, mitotic microglia, mechanical allodynia, and heat hyperalgesia. **Figure 4-figure supplement 1.** Stratagem for generating *Mtor-cKO^MG^* mice.

### *Mtor* ablation in DRG neurons suppressed elevation of subsets of nerve injury- induced genes

To determine the downstream molecular targets of mTOR in DRG neurons involved in neuropathic pain, we performed RNA sequencing of DRGs from *Mtor^fl/fl^* and *Mtor-cKO^Adv^* mice before and 7 days after SNI surgery. In total, the expression levels of 189 genes (155 upregulated and 34 downregulated), were significantly changed (by at least two folds, *p*<0.05) in the injured DRGs 7 days after SNI in *Mtor^fl/fl^* mice (**Figure 5A- C**). A large number of the upregulated genes, including those associated with injury (*Activating transcription factor 3*, *Atf3* and *Small proline-rich protein 1A*, *Sprr1a*), G-protein coupled receptors (*Gpcrs*, including *Gpr151* and *Gpr119*), neuropeptides (*Npy*, *Gal*, and *Nts*), cytokines (*Colony stimulating factor 1*, C*sf1* and *Interleukin 1b*, *Il1b*) have been previously reported in response to nerve injury (**Figure 5B**) (Wu et al., 2016; Reinhold et al., 2015; Guan et al., 2016; Peng et al., 2016), verifying the reliability of the RNA-seq data. Gene ontology analysis demonstrated that injury-affected genes were primarily enriched in four molecular functions (**Figure 5C**), including receptor ligand activity, hormone activity and neuropeptide receptor binding and activity.

**Figure 5.**
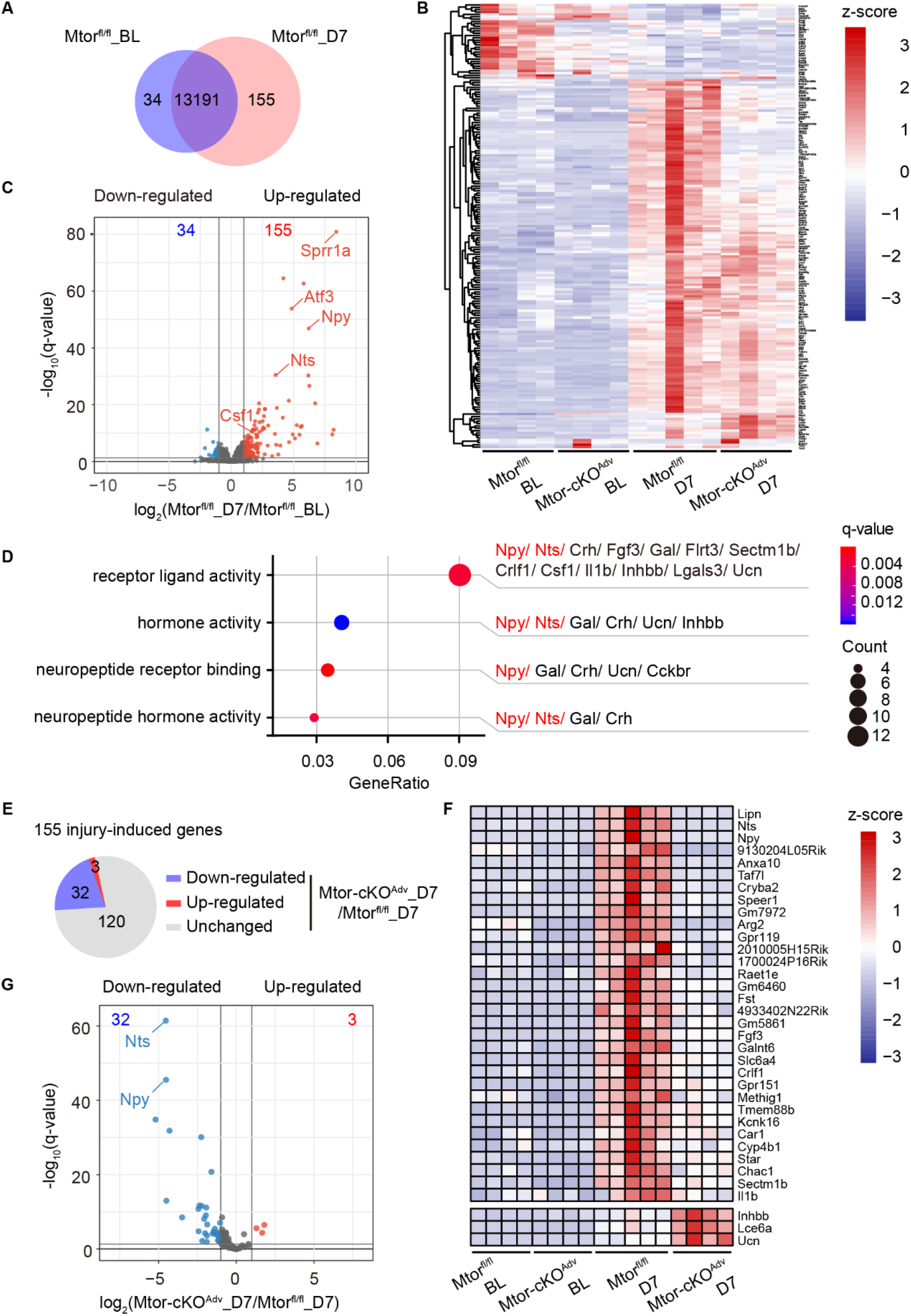
Ablation of *Mtor* in DRG neurons suppressed elevation of nerve injury-induced genes. (**A**) Venn diagram of DEGs identified in DRG before and after SNI (day 7) in *Mtor^fl/fl^* mice (155 upregulated and 34 downregulated). (**B**) Heat map of 189 DEGs by hierarchical clustering using z-score values (*n* = 4-5 mice per group). (**C**) Volcano plot of DRG transcripts before and after SNI (day 7) in *Mtor^fl/fl^* mice. Red dots indicate 155 upregulated genes and blue dots indicate 34 downregulated genes after SNI. (**D**) GO analysis of 155 upregulated genes after SNI and regroup into molecular function terms. All genes in each term are listed. (**E**) Pie chart of 155 injury-induced genes with 32 downregulated and 3 upregulated in *Mtor-cKO^Adv^* mice after SNI. (**F**) Heat map of 35 DEGs in all samples using Z-score values. Only 3 (*Inhbb, Lce6a and Ucn*) of the 155 injury-induced genes are upregulated upon deletion of *Mtor* in DRG neurons. (**G**) Volcano plot of 35 DEGs in control and *Mtor-cKO^Adv^* mice after SNI. Red dots indicate 3 upregulated genes and blue dots indicate 32 downregulated genes after mTOR ablation. BL, baseline; D, day; DEGs, differentially expressed genes. **Figure 5-source data 1.** FPKMs of differentially expressed genes (189 genes) used to generate Figure B. **Figure 5- source data 2.** FPKMs of differentially expressed genes (35 genes) upon *Mtor* ablation used to generate Figure F. **Figure 5-figure supplement 1.** Quantitative RT-PCR of downregulated DEGs identified in RNA sequencing.

Importantly, approximately 1/5 (32 in 155 genes) of injury-induced genes were suppressed after mTOR ablation (**Figure 5E**). In particular, the expression of two neuropeptide genes *Npy* and *Nts*, induced by approximately 73.5 and 11.7 folds after injury, was strikingly reduced to 3.75 and 0.57 folds after ablation of *Mtor* in DRG neurons. By contrast, expression of another two injury-induced neuropeptide genes, such as *Corticotropin releasing hormone* (*Crh*) and *Gal* remained largely unaffected, suggesting that mTOR specifically regulates the expression of subsets of injury-responsive genes (**Figure 5E-G**). The reduced expression of *Npy, Nts*, and other genes (as indicated) in *Mtor-cKO^Adv^* mice was further verified by qRT-PCR analysis (**Figure 5-figure supplement 1**). Notably, while mTOR was transiently activated during the first week after nerve injury, it may have long-term impacts on downstream molecules. Collectively, these data demonstrate that mTOR regulates the transcription of a number of injury-induced genes.

### Injury-activated mTOR is required for NPY induction in DRG neurons

NPY is widely distributed in the central and peripheral nervous system (Allen et al., 1983). It is absent in DRG neurons under homeostatic conditions but dramatically upregulated after peripheral nerve injury (Wakisaka et al., 1991; Xiao et al., 2002). However, nothing is known about the mechanisms regulating NPY induction after nerve injury. We also observed prominent induction of *Npy* in DRG neurons after nerve injury, which lasted for at least 4 weeks with gradually reduced levels after day 14 (**Figure 6A and B**). Immunofluorescence analysis revealed that 94.2% of NPY^+^ neurons were co-labelled with ATF3, a marker for neuronal injury marker (**Figure 6C and D**). Moreover, 89.6% of NPY^+^ neurons expressed p-S6 (**Figure 6E and F**), and co-localized with NF160/200 in the injured DRGs (**Figure 6-figure supplement 1**), suggesting that NPY is selectively induced in injured large-sized mechanoreceptors with mTOR activation. It is noteworthy that *Mtor* ablation nearly eliminated NPY induction (**Figure 6E and G**), indicating that mTOR inactivation suppressed NPY transcription.

**Figure 6.**
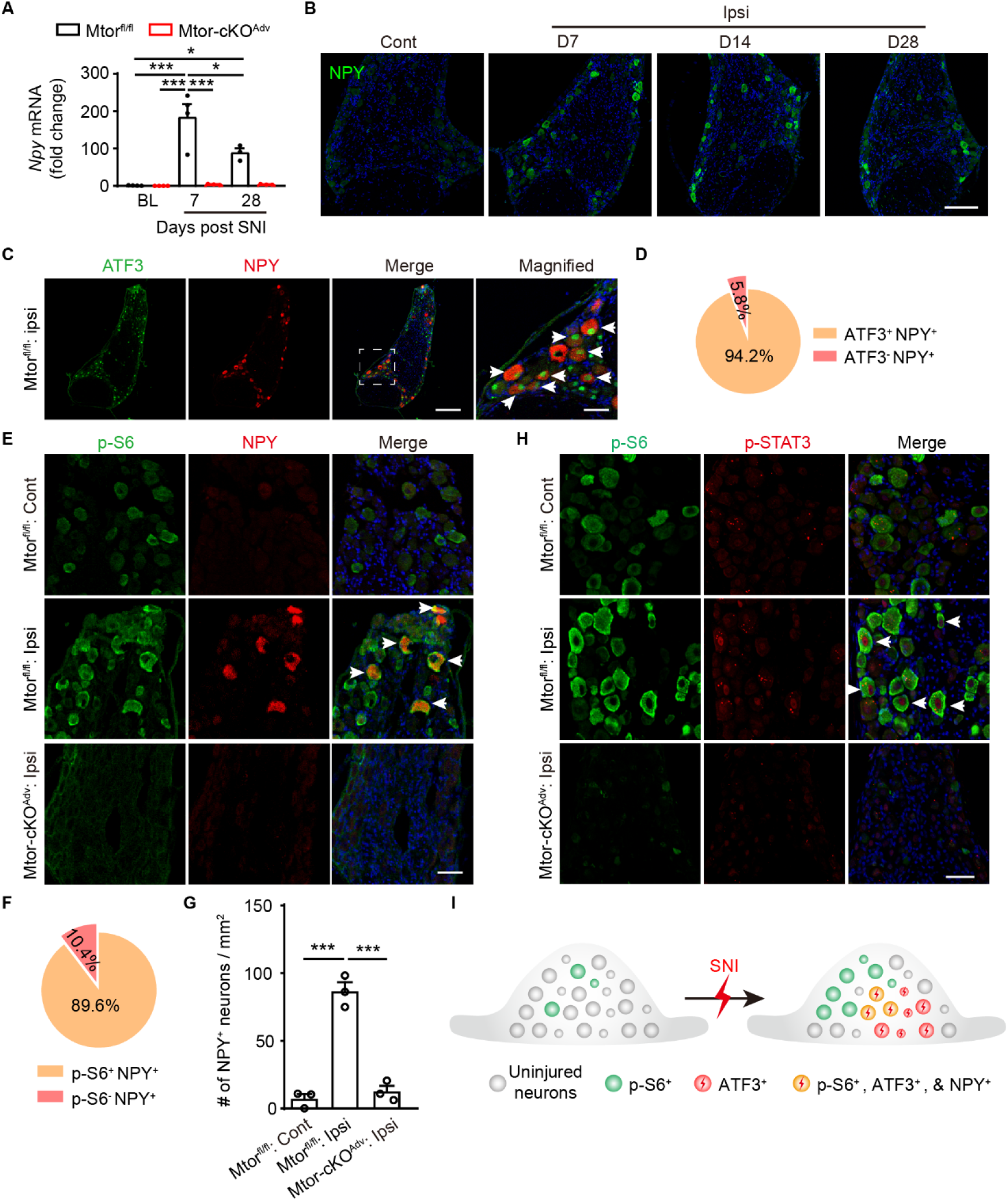
Activation of mTOR is required for NPY induction in DRG neurons after SNI. (**A**) Quantitative RT-PCR of *Npy* transcripts in the ipsilateral DRG from both *Mtor^fl/fl^* and *Mtor-cKO^Adv^* mice at indicated time points after SNI (*n* = 3-4 mice per time point). (**B**) Representative images of NPY staining in DRG from *Mtor^fl/fl^* mice at indicated times. Scale bars, 100 μm. (**C**) Representative images of ATF3 and NPY staining in the ipsilateral DRG from *Mtor^fl/fl^* mice at day 7 after SNI. Arrows indicate ATF3^+^ NPY^+^ neurons. Dotted boxes show regions of higher magnification in the DRG. Scale bars, 200 μm for low magnification images and 50 μm for high magnification images. (**D**) A pie chart showing the ratio of NPY^+^ in total ATF3^+^ neurons in the ipsilateral DRG from *Mtor^fl/fl^* mice at day 7 after SNI. (**E**) Representative images of NPY and p-S6 staining in contralateral or ipsilateral DRG at day 7 after SNI in *Mtor^fl/fl^* and *Mtor-cKO^Adv^* mice. Arrows indicate co-labeled neurons. Dotted boxes show regions of higher magnification in the DRG. Scale bar, 50 μm. (**F**) A pie chart indicating the ratio of NPY^+^ neurons in all p-S6^+^ neurons in ipsilateral DRG from *Mtor^fl/fl^* mice at day 7 after SNI. (**G**) Quantification of NPY^+^ neurons in *Mtor^fl/fl^* and *Mtor-cKO^Adv^* mice at day 7 after SNI (*n* = 3 mice per group). (**H**) Representative images of p-S6 and p-STAT3 staining in contralateral or ipsilateral DRG at day 3 after SNI in *mTOR^fl/fl^* and *Mtor-cKO^Adv^* mice. Scale bar, 50 μm. (**I**) Schematic diagram demonstrating that NPY is selectively induced in p-S6^+^ and ATF3^+^ injured neurons in ipsilateral DRG. Values are means ± SEM. * *p*<0.05, and *** *p*<0.001, one-way ANOVA followed by Bonferroni’s *post hoc* tests among groups. Cont, contralateral; Ipsi, ipsilateral. **Figure 6-source data 1.** Raw data of *Npy* transcripts and number of NPY^+^ neurons. **Figure 6-figure supplement 1.** NPY was expressed in large-sized mechanoreceptors.

Previous studies indicated that the promoter regions of *Npy* and *Nts* genes harbor signal transducer and activator of transcription 3 (STAT3)-binding site-like elements and that dominant negative expression of STAT3 attenuated leptin-induced *Npy* and *Nts* expression (Muraoka et al., 2003; Cui et al., 2005). In addition, activated mTOR has been shown to phosphorylate STAT3 to promote its nuclear entry and gene transcription (Laplante and Sabatini, 2013). Importantly, a recent study showed that phosphorylation of STAT3 was elevated in DRG neurons after nerve injury (Chen et al., 2016). We therefore tested whether phosphorylation of STAT3 might be involved in mTOR-mediated NPY induction. We found that nerve injury-increased STAT3 phosphorylation was completely blocked after mTOR ablation, suggesting a correlation between STAT3 phosphorylation and NPY induction (**Fig. 6H**). Together, these data demonstrate that activation of mTOR is required for nerve injury-induced NPY elevation in DRG neurons (**Figure 6I**).

### Nerve injury-induced NPY enhances nociceptor excitability

NPY has been shown to increase the excitability of DRG neurons (Abdulla and Smith, 1999a). To examine whether mTOR-promoted NPY induction enhances the excitability of nociceptors, we carried out electrophysiological recording of small-sized nociceptors at 7 days SNI. As expected, nociceptors from *Mtor^fl/fl^* mice displayed increased number of action potentials and lower rheobase 7 days after SNI (**Figure 7**). By contrast, the number of spikes was significantly reduced in mTOR-deficient neurons after injury. However, incubation of NPY with mTOR-deficient neurons significantly restored the number of action potentials and reduced rheobase, suggesting that NPY loss contributes to the reduced nociceptor excitability in the absence of mTOR.

**Figure 7.**
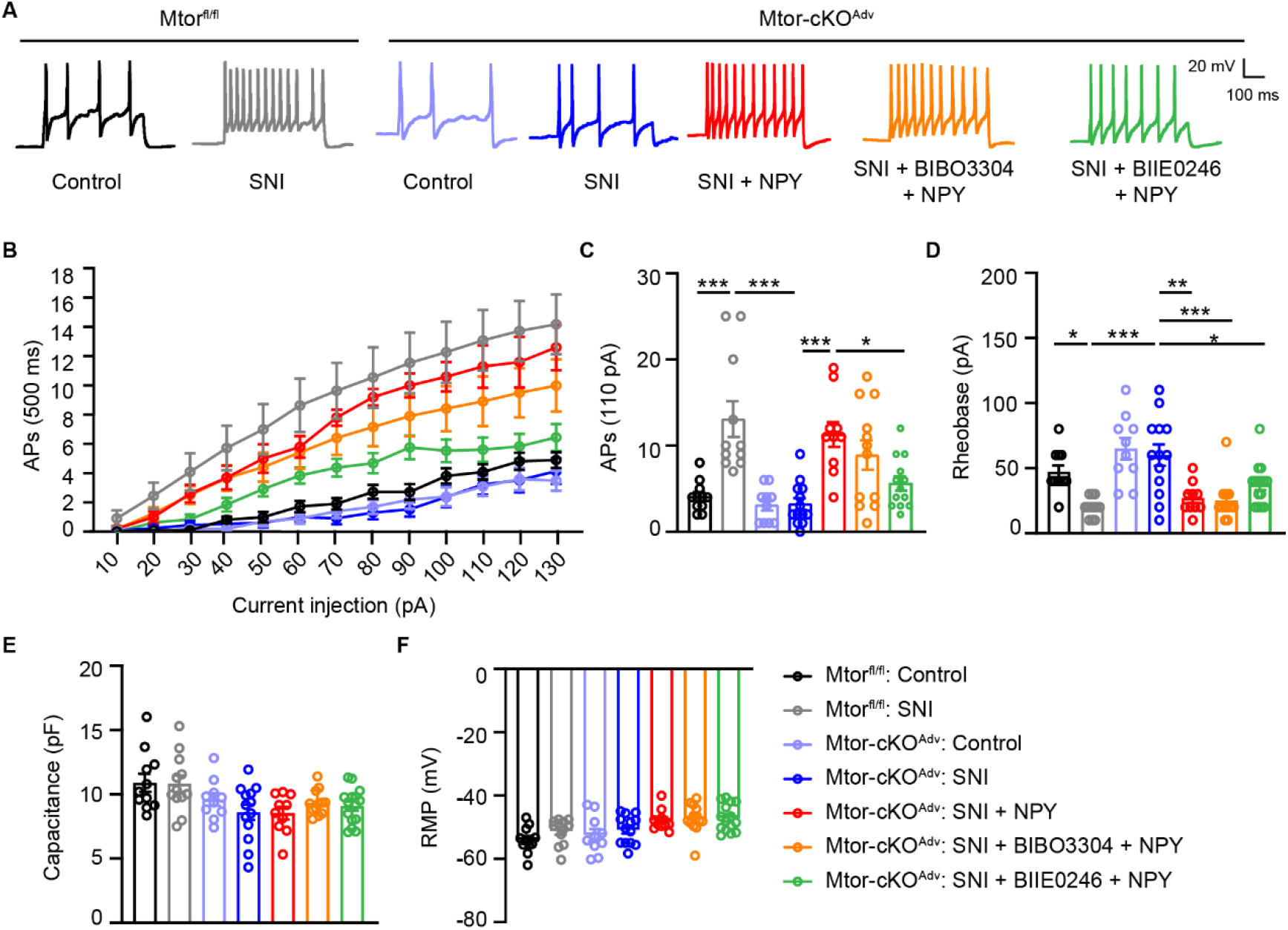
NPY enhances nociceptor excitability through Y2R. (**A**) Representative AP traces elicited by intracellular injection of 110 pA depolarizing currents on dissociated DRG neurons from resting membrane potentials (RMP) in *Mtor^fl/fl^* mice and *Mtor-cKO^Adv^* mice with or without SNI. NPY (300 nM), BIBO3304 (1 μM) and BIIE0246 (1 μM) are replenished in medium as indicated. (**B**) The response of *Mtor^fl/fl^* and *Mtor-cKO^Adv^* DRG neurons across a series of 500 ms depolarizing current pulses in 10 pA increment from 0 pA to 130 pA, in the presence or absence of NPY, BIBO3304 or BIIE0246 (*n* = 10-13 neurons per group). (**C**) Quantification of APs evoked by input current at 110 pA (*n* = 10-13 neurons per group). (**D**) Averaged values of rheobase currents in DRG neurons among groups measured in I-clamp (*n* = 10-13 neurons per group). (**E-F**) Quantification of membrane capacitance (**E**) and RMP (**F**) among groups (*n* = 10-13 neurons per group). BIBO3304, Y1R antagonist; BIIE0246, Y2R antagonist. Values are means ± SEM. * *p*<0.05, ** *p*<0.01, and *** *p*<0.001, one-way ANOVA followed by Bonferroni’s *post hoc* tests among groups. AP, action potential; RMP, resting membrane potentials. **Figure 7-source data 1.** Raw data of APs, rheobase currents, membrane capacitance, and RMP. **Figure 7-figure supplement 1.** Distinct expression pattern of NPY (*) and Y2R (arrows) by immunofluorescence analysis.

Studies have demonstrated different responses of DRG neurons to different NPY receptor agonists (Wiley et al., 1993; Abdulla and Smith, 1999b; Abdulla and Smith, 1999a). For example, Y2R agonists increased neuronal excitability of small DRG neurons, whereas Y1R agonists barely showed any effects (Abdulla and Smith, 1999a). We verified the distinct expression pattern of NPY and Y2R on large-sized mechanoreceptors and small-sized nociceptors by immunofluorescence analysis (**Figure 7-figure supplement 1**) (Brumovsky et al., 2005). To determine which receptor mediates NPY-elicited excitatory effects, Y1R or Y2R antagonist was incubated with DRG neurons for 30 min before NPY addition. We found that blocking Y2R but not Y1R activity substantially reduced the number of action potentials after NPY addition, suggesting that Y2R mediates NPY-induced elevated excitation (**Figure 7A-C**).

### Peripheral NPY replenishment reversed analgesic effects of *Mtor* ablation through Y2R

NPY has been shown to elicit biphasic effects in pain processing by binding to different receptors in DRG or spinal neurons (Brumovsky et al., 2007). Given that mTOR ablation simultaneously delayed pain onset and suppressed NPY induction, we tested whether mTOR inactivation alleviated pain via NPY loss. We first administered a small dose of NPY (0.2 nmol), as previously suggested (Tracey et al., 1995; Brumovsky et al., 2005), into the hind paw of normal mice and observed prominent mechanical allodynia and heat hyperalgesia approximately 30 minutes after injection, supporting the pro-nociceptive effects of peripheral NPY (**Figure 8A-C**). By injecting NPY into the ipsilateral hind paw of *Mtor-cKO^Adv^* mice, we observed robust mechanical allodynia, and to a lesser extent, heat hyperalgesia in *Mtor-cKO^Adv^* mice (**Figure 8E and F**). Moreover, blocking Y2R, rather than Y1R, before NPY administration substantially reduced NPY-induced mechanical allodynia (**Figure 8E**), further supporting the role of Y2R in mediating NPY-elicited pro-nociceptive effects. Collectively, our data demonstrate that mTOR-induced NPY production in DRG neurons is essential for the development of neuropathic pain via Y2R-mediated signaling.

**Figure 8.**
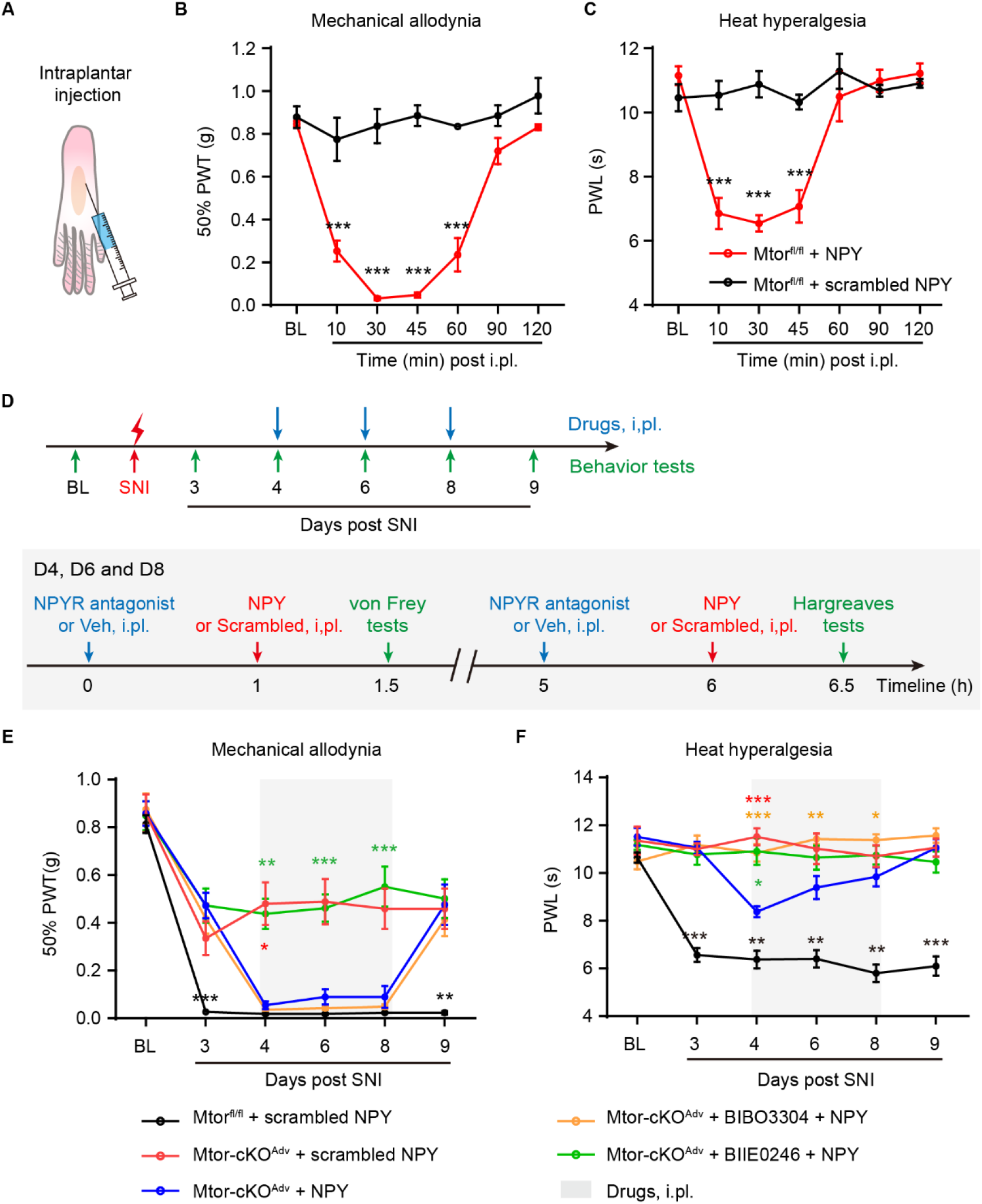
Intraplantar injection of NPY reversed analgesic effects of *Mtor* ablation through Y2R. (**A**) Schematic diagram indicating intraplantar (i.pl.) injection. (**B-C**) NPY (0.2 nmol) i.pl. injection into normal *Mtor^fl/fl^* mice hind paw leads to transient mechanical allodynia (**B**) and heat hyperalgesia (**C**) within an hour (*n* = 4 mice per group). (**D**) Experimental schedule showing the timeline of i.pl. injection of drugs (including NPY, NPYR antagonist and vehicle) and behavior tests. Behavior tests were measured before and after SNI as indicated. Drugs were injected at day 4, 6 and 8. (**E-F**) Measurement of mechanical allodynia (**E**) and heat hyperalgesia (**F**) in Mtorfl/fl and Mtor-cKOAdv mice with i.pl. injection with NPY (0.2 nmol), scrambled NPY (0.2 nmol), BIBO3304 (5 nmol) or BIIE0246 (50 nmol) at day 4, 6, and 8 after SNI (n = 6- 11 mice per group). BIBO3304, Y1R antagonist; BIIE0246, Y2R antagonist. Values are means ± SEM. * *p*<0.05, ** *p*<0.01, and *** *p*<0.001 *vs. Mtor-cKO^Adv^* with NPY, two-way ANOVA followed by Bonferroni’s *post hoc* tests among groups. BL, baseline; i.pl., intraplantar; Veh, vehicle; PWT, paw withdraw threshold; PWL, paw withdraw latency. **Figure 8-source data 1.** Raw data of mechanical allodynia and heat hyperalgesia.

## DISCUSSION

Neuropathic pain is a maladaptive response of the nociceptive pathway to the nerve injury. Both peripheral and central sensitization have been shown to contribute to the persistent pain (Colloca et al., 2017). Peripheral nociceptor sensitization is a key trigger in neuropathic pain, as inhibiting nociceptor activity by anesthetics effectively blocks pain (Colloca et al., 2017). In this study, we uncover a previously unrecognized mechanism, by which injury-induced mTOR activation and downstream transcriptional factor STAT3 drives NPY synthesis to enhance nociceptor excitability and promote pain development through Y2R. Considering the distinct distribution patterns of NPY and Y2R in large-sized mechanoreceptors and small-sized nociceptors, mTOR-driven pain may involve an intra-ganglia communication between NPY-expressing mechanoreceptors and Y2R-expressing nociceptors.

Basal levels of mTOR activity are present in a small subset of large-sized myelinated sensory neurons in naïve mice (Lisi et al., 2015; Geranton et al., 2009). In the present study, we observed increased mTOR activation predominantly occurs in large sensory neurons and spinal microglia after nerve injury. While pharmacologically blocking mTOR activity has raised controversies regarding its role in pain (Obara et al., 2011; Geranton et al., 2009; Melemedjian et al., 2013; Khoutorsky et al., 2015), we found that selective ablation of mTOR in primary sensory neurons, which disrupted both mTORC1 and mTORC2, robustly prevented the early onset of nerve injury-triggered allodynia and heat hyperalgesia for 2 weeks. Selective ablation of Raptor or Rictor in DRG neurons are needed to distinguish roles of mTORC1 and mTORC2 signaling in neuropathic pain (Laplante and Sabatini, 2012). In contrast to the ERK activation following mTORC1 inhibition as previously described (Melemedjian et al., 2013), we did not observe ERK activation in DRG neurons (data not shown) after genetic inactivation of mTOR in SNI models. The delayed onset of mechanical allodynia after mTOR ablation are in line with the temporal activation of mTOR and expression of downstream effectors after nerve injury, emphasizing a central role of mTOR in promoting neuropathic pain development. Additional maladaptive changes other than mTOR signaling may contribute to the late-phase pain.

Nerve injury-induced *de novo* synthesis of a large number of molecules are implicated in the hypersensitive nociception (Wang et al., 2021; Zhao et al., 2017). For example, injury-induced CSF1 in DRG neurons, a cytokine required for microglial and macrophage expansion, has recently been shown to contribute to mechanical hypersensitivity (Yu et al., 2020; Guan et al., 2016; Peng et al., 2016). Also, removal of the eukaryotic initiation factor 4E-binding protein 1 (4E-BP1), a negative regulator of protein translation downstream of mTOR, induced pain hypersensitivity through enhanced translation of neuroligin 1 even in the absence of nerve injury, further stressing the importance of mTOR-mediated protein synthesis in pain hypersensitivity (Khoutorsky and Price, 2018; Khoutorsky et al., 2015; Yousuf et al., 2020). Our findings that mTOR is required for nerve injury-induced *Npy* and *Nts* transcription demonstrate novel links between mTOR activation to neuropeptide production.

Intriguingly, as a serine/threonine kinase that is primarily engaged in translational control, mTOR is unlikely to directly promote *Npy* or *Nts* transcription. In search for potential mTOR-regulated transcriptional factors upstream of *Npy* or *Nts* genes, we observed suppressed phosphorylation of STAT3, but not C-Jun or CREB (data not shown), in DRG neurons after mTOR deletion. Since STATs-like binding elements are present in the promoter region of *Npy* gene (Muraoka et al., 2003), it is likely that activated mTOR induces *Npy* transcription by phosphorylating STAT3, thereby promoting STAT3 nuclear entry and downstream gene transcription. While previous studies primarily suggested that mTOR contributes to pain sensitivity through translational control (Khoutorsky and Price, 2018; Melemedjian and Khoutorsky, 2015), our study demonstrate a non-translational mechanism of mTOR involving STAT3- NPY production in pain regulation.

Nerve injury often induces nociceptor hyper-excitability to provoke pain hypersensitivity. However, this hyper-excitability was lost after ablation of mTOR in DRG neurons, along with elimination of NPY. NPY has been shown to elicit both anti- nociceptive and pro-nociceptive effects, depending on the subtypes of its receptors in the central and peripheral nervous system (Brumovsky et al., 2007; Diaz-delCastillo et al., 2018). We found that NPY was selectively induced in injured large-sized sensory neurons, suggesting the peripheral effects of NPY. In contrast to previous studies showing that NPY triggers analgesia by inhibiting superficial dorsal horn interneurons through Y1R (Taiwo and Taylor, 2002; Miyakawa et al., 2005; Nelson and Taylor, 2021), we observed that peripheral administration of NPY promotes pain via Y2R. Replenishing NPY enhanced nociceptor excitability, while peripherally blocking Y2R, but not Y1R, prevented these effects, suggesting that mTOR drives NPY production to enhance nociceptor excitability through Y2R. Consistent with our observations, a previous study indicated that Y1R agonist had no effect on small DRG neurons, whereas Y2R agonist enhanced neuronal excitability (Abdulla and Smith, 1999a). A reasonable explanation is that Y2R attenuated calcium-sensitive potassium- conductance, thereby inducing nociceptor depolarization and excitability (Abdulla and Smith, 1999a; Abdulla and Smith, 1999b).

It is noteworthy that NPY elevation is exclusively observed in large-diameter mechanoreceptors, whereas Y2R is predominantly distributed in small-diameter nociceptors (Brumovsky et al., 2005). The distinct but adjacent distribution of NPY suggests a paracrine ‘somatic cross excitation’ model within DRGs, by which NPY released from the large-diameter injured neurons acts on neighboring small-diameter Y2R-expressing neurons (Brumovsky et al., 2007). Through intra-ganglionic transmission (Brumovsky et al., 2007), NPY signals derived from large injured mechanoreceptors are able to sensitize Y2R-expressing nociceptors, thereby contributing to mechanical allodynia. In line with this concept, blocking Y2R effectively alleviated mechanical allodynia. Our findings thus provide an important mechanism for mechanical allodynia engaging mTOR-driven NPY-Y2R communication between mechanoreceptors and nociceptors in neuropathic pain. Other than NPY, mTOR-driven expression of NTS and GPCRs likely coordinately contribute to the full development of neuropathic pain.

Microglia activation in SDH have been shown to contribute to neuropathic pain (Inoue and Tsuda, 2018). Moreover, mTOR-mediated metabolic reprogramming are required for induction of inflammatory factors and cytokines in microglia (Hu et al., 2019), which indicated that mTOR activation in microglia may be involved in neuropathic pain. We found that *Mtor* deletion in microglia reduced microgliosis; however, it did not have significant effects on neuropathic pain. This is likely due to the fact that mTOR was activated in less than 50% of microglia in the SDH and that mTOR ablation only partially reduced microgliosis, which might be insufficient to inhibit pain development. Consistent with this notion, removal of microglia only ameliorated mechanical allodynia during the first 3 days after nerve injury, whereas removal of both microglia and peripheral monocytes/macrophage prevented neuropathic pain development (Peng et al., 2016).

In summary, we demonstrate that nerve injury-induced aberrant mTOR activation in sensory neurons promotes pain development. While mTOR has been shown to affect the expression or function of hundreds of molecules, the present study is the first that links mTOR to NPY signaling in sensitizing nociceptive pathway to drive neuropathic pain. As mTOR inhibitors are in clinical use and Y2R receptor antagonists are readily available, our findings also provide new perspectives for clinically treating neuropathic pain by peripherally modulating mTOR and NPY-Y2R signaling.

## Materials and Methods

### Key resources table

**Table 1.**
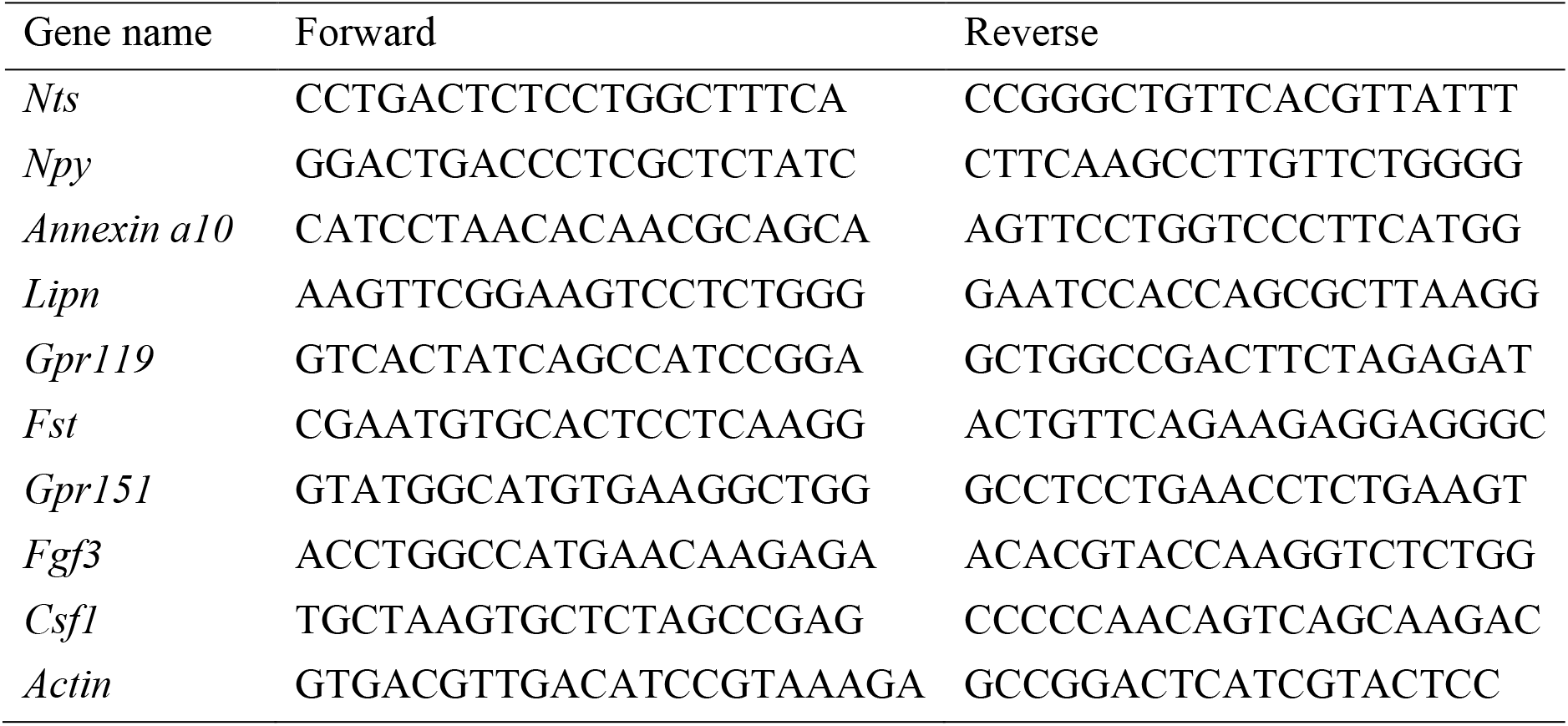
Primer sequences in RT-PCR.

### Animals

Adult male mice (8-12 weeks) were used for biochemical and behavioral tests, and young mice (4-6 weeks) for whole-cell patch clamp recording. C57BL/6J mice were purchased from Shanghai Slac Laboratory Animal Corporation (China). *Cx3cr1^EGFP/+^*, *Cx3cr1^creER/+^*, *Mtor floxed* (*Mtor^fl/fl^*) and *Advillin^cre^* (*Adv^cre^*) mice with C57BL/6J background were purchased from the Jackson Laboratory (ME, USA). All animals were housed under a 12-h light/dark cycle with food and water available. To selective knockout the *Mtor* gene in microglia, mice bearing the floxed allele of the *Mtor* gene (*Mtor^fl/fl^*) were crossed with *Cx3cr1^creER/+^*mice. *Cx3cr1^creER/+^::Mtor^fl/fl^* mice received two doses of 10 mg tamoxifen citrate (TAM, Meilunbio, China) or vehicle in 48-hour intervals. TAM induced the expression of Cre recombinase in both resident microglia and peripheral monocytes. Since monocytes have a rapid turnover rate, Cre expression is eliminated in peripheral monocytes but maintained in resident microglia 4-6 weeks after TAM induction (Parkhurst et al., 2013), thus allowing selective deletion of *Mtor* in microglia (*Mtor-cKO^MG^*) but not in monocytes. Control mice were *Cx3cr1^creER/+^::Mtor^fl/fl^* littermates without TAM induction and *Cx3cr1^creER/+^* mice with TAM induction. For selective ablation of *Mtor* in DRG sensory neurons, *Mtor^fl/fl^* mice were crossed with *Adv^cre^* mice to obtain the *Adv^cre^::Mtor^fl/fl^* (*Mtor-cKO^Adv^*) mice. *Mtor- cKO^Adv^* mice enabled *Mtor* deletion in DRG neurons but leave spinal cord unaffected. Control mice were *Mtor^fl/fl^* littermates without Cre promotor.

### Cre-mediated recombination of the *Mtor^flox^* allele

Primers used for analyses of *Mtor* floxed alleles were as the following: *Mtor-P1* (5’- GCTCTTGAGGCAAATGCCACTATCACC-3’), *Mtor-P2* (5’- TCATTACCTTCTCATCAGCCAGCAGTT-3’), *Mtor-P3* (5’-TTCATTCCCTTGAAAGCCAGTCTCACC-3’). Primer pair P1/P2 was used for genotyping floxed mTOR alleles that generated a 480 bp DNA fragment in PCR (Risson et al., 2009). Upon Cre- mediated recombination, *P1/P3* pair produced a recombined *Mtor* gene fragment of 520 bp with excision of exons 1-5 (**Figure 4-figure supplement 1**) (Risson et al., 2009).

### Neuropathic pain model

Spared nerve injury (SNI) models were used to induce neuropathic pain as previously described (Decosterd and Woolf, 2000). Mice were anesthetized with sodium pentobarbital (100 mg/kg) intraperitoneally. The left hindlimb was shaved, and the skin was disinfected with iodophor. After blunt separation of biceps femoris muscle, 3 distal branches of sciatic nerve were exposed and the tibial and common peroneal nerves were ligated with 5-0 silk sutures, with care to avoid injury to the sural nerve. The ligated branches were then transected distal to the ligature and a 2-3 mm distal nerve stump was removed. To minimize the number of animals used in the experiments, the right hindlimb was performed with a sham surgery after sciatic nerve exposure without nerve ligation and transection. To analyze NPY transport from the DRG to the spinal cord, we ligated the ipsilateral L4 central axonal branches immediately after SNI. After the surgery, the incision was closed using 5-0 silk sutures. The injured side was then regarded as the ipsilateral side, and the uninjured as the contralateral one.

### Western Blotting

Bilateral lumbar 4 and 5 (L4-L5) DRGs and dorsal horns of L4-L5 spinal cord were isolated at certain time points after SNI surgery, snap-frozen in liquid nitrogen and stored at -80℃. Tissues were homogenized in RIPA lysis buffer (Beyotime, China) with protease inhibitor (Cat# S8830, Sigma-Aldrich, MO, USA) and phosphatase inhibitor (Cat# A32961, Thermo Fisher, MA, USA) using ultrasonic cell disruptor. The homogenates were centrifuged at 4 ℃ for 30 minutes at 10,000 g and the supernatants were collected. Proteins were separated by 10% SDS-polyacrylamide gels and transferred to polyvinylidene difluoride membranes (Millipore, Germany), followed by blocking, primary antibodies and horseradish peroxidase (HRP)-conjugated secondary antibodies (1:10000, Jackson ImmunoResearch, PA, USA) incubation. The proteins were detected using enhanced chemiluminescence regents (ECL, Amersham Pharmacia Biotech, NJ, USA) according to the manual.

### Immunofluorescence analysis

After deeply anaesthetized with sodium pentobarbital, mice were perfused with saline and subsequently 4% paraformaldehyde (PFA, Sigma-Aldrich). The spinal cord and L4-L5 DRGs were dissected, post-fixed in 4% PFA, and transferred to 30% sucrose in 0.1 M phosphate buffer (pH=7.2) for 2 days. Samples were embedded in optimal cutting temperature (OCT) and transverse sections were cut using freezing microtome (Lecia Biosystems, Germany) at a thickness of 15 μm. To label Isolectin B4 (IB4)-positive neurons in DRG, slices were blocked with 10% (wt/vol) normal bovine serum albumin (BSA) for 1 hour at room temperature, and incubated with 1 μg/mL IB4 diluted in phosphate buffered saline (PBS) at room temperature for 2 hours. Sections were washed with Tris buffered saline (TBS) and then incubated with anti-p-S6 (1:1000) antibody. For staining with other antibodies, sections were antigen-retrieved in citrate buffer (10 mM sodium citrate, 0.05% Tween-20, pH 6.0) or Tris-EDTA (10 mM Tris, 1 mM EDTA, 0.05% Tween-20, pH 9.0) as appropriate at 95 ℃ for 20 minutes and permeabilized with 0.5% Triton X-100 for 10 minutes at room temperature. After blocked with 10% (wt/vol) BSA, sections were incubated overnight at 4 °C with following primary antibodies: rabbit anti-p-S6 (1:1000), mouse anti-p-S6 (1:2000 - 1:4000), mouse anti-NeuN (1:1000), mouse anti-NF160/200 (1:2000), mouse anti- CGRP (1:1000), rat anti-BrdU (1:800), and goat anti-GFP (1:1000), rabbit anti-Iba1 (1:800), rabbit anti-NPY (1:1000), mouse anti-GFAP (1:800), mouse anti-ATF3 (1:200), rabbit anti-p-STAT3 (Ser727) (1:500). Sections were then washed in TBS with 0.5% tween (TBS-T) and incubated with appropriate secondary antibodies (1:1000) for 1.5 hours at room temperature. For NPY and Y2R staining, since both anti-NPY and Y2R antibodies were raised in rabbits, the multiple-color immunochemistry kit (Cat# abs50012, Absin, China) was used following the manufacturer’s instructions. The specificity of the staining using this kit was first validated by double staining of rabbit anti-NPY and Iba1 antibodies that showed no overlaps. Following the anti-NPY incubation, rabbit horseradish peroxidase (HRP)-conjugated secondary antibody (1:1000) was applied and incubated for 1.5 hours. Sections were than washed in TBS- T and incubated with Tyramide Signal Amplification (TSA) reagent for 10 minutes. Antibody eluent (Cat# abs994, Absin) was used to wash out anti-NPY and HRP- conjugated antibody. After washing, sections were incubated with anti-Y2R antibody (1:500) and followed by incubation with appropriate secondary antibodies (1:1000) according to species of the first antibody. DAPI (Beyotime) was used to label cell nuclei in tissue sections. The immunofluorescence images were captured by FV-1200 confocal microscope (Olympus, Japan). The density or percentage of positive cells in SDH and DRGs were counted and calculated using 3 sections from each animal. Mean intensity of interested regions were evaluated using *ImageJ* software.

### Drug administration

BrdU was used to label proliferating cells in the spinal cord after the SNI surgery. The BrdU labeling procedure was carried out as described before (Gu et al., 2016), with two intraperitoneal injections (100 mg/kg) daily one day before the surgery until 7 days post-surgery. For intraperitoneal treatment of rapamycin, mice were administrated with rapamycin (5 mg/kg) or vehicle daily one day before SNI until 7 days post-surgery. For local intraplantar (i.pl.) injection, drugs (0.2 nmol NPY, 0.2 nmol scrambled NPY, 5 nmol BIBO3304 trifluoroacetate or 50 nmol BIIE0246) in 20 μl saline were injected using a syringe with a 30-gauge needle. Dosages of NPY and its antagonists were referred to the previous studies (Tracey et al., 1995; Sapunar et al., 2011). NPY receptor antagonists were injected 1 hour before NPY injection. To assess the effects after i.pl. injection, behavioral tests were finished in 30-40 minutes after NPY or scrambled peptide injection. The von Frey and Hargreaves tests were used for an interval of at least 4 hours.

### RNA sequencing

Bilateral SNI were performed in *Mtor^fl/fl^* and *Mtor-cKO^Adv^* mice to minimize the animals used in the experiment. In total, 4 lumber DRGs (bilateral L4 and L5 DRGs) were collected from each mouse before or 7 days after SNI. RNAs were isolated using RNeasy micro kit (Cat# 74004, QIAGEN, Germany) according to the manufacturer’s instructions. RNA sequencing (RNA-seq) libraries were constructed and sequenced by BGISEQ-500 (BGI, China). After quality control, the raw RNA-seq data were filtered to obtain the clean data used for alignment to the mouse genome (Mus musculus GRCm38.p5, NCBI). Based on these read counts, normalization and differential gene expression were performed using DESeq2 on *R* (version 3.5.3). Genes with fragments per kilobase million (FPKM) lower than 1 (FPKM<1) in all groups were excluded from the subsequent analyses. Statistical significance of differentially expressed genes (DEGs) was calculated based on the raw counts of individual genes, with an absolute fold change greater than 2 and adjusted p-value (q-value) less than 0.05.

Volcano plots and heatmaps were visualized by R (the ggplot2 and gplots packages, respectively). Gene Ontology (GO) enrichment in the molecular function category were visualized by R (bioconductor package “org.Hs.eg.db” and “cluster profiler” package).

### Quantitative RT-PCR

Total RNA from DRG was extracted using RNeasy micro kit and reverse-transcribed using PrimeScript RT Reagent Kit (Cat# RR037A, TaKaRa, Japan). Real-time PCR was performed using the SYBR Premix Ex Taq™ (Cat# DRR041A, Takara) on a LightCycler 480 Instrument II Real-Time PCR Detection System (Roche). Primer sequences are provided in the **Table 1**. The relative expression was measured using the 2^−ΔΔCt^ method. Briefly, the threshold cycle (Ct) values of target genes were determined automatically by LightCycler 480 II software. ΔCt = Ct_(Target genes)_ – Ct_β-actin_. ΔΔCt = ΔCt_(Target genes)_ − ΔCt_(average ΔCt of control)_. Relative fold changes were determined by 2^−ΔΔCt^ and normalized to the expression levels of *Actin (Livak and Schmittgen, 2001)*.

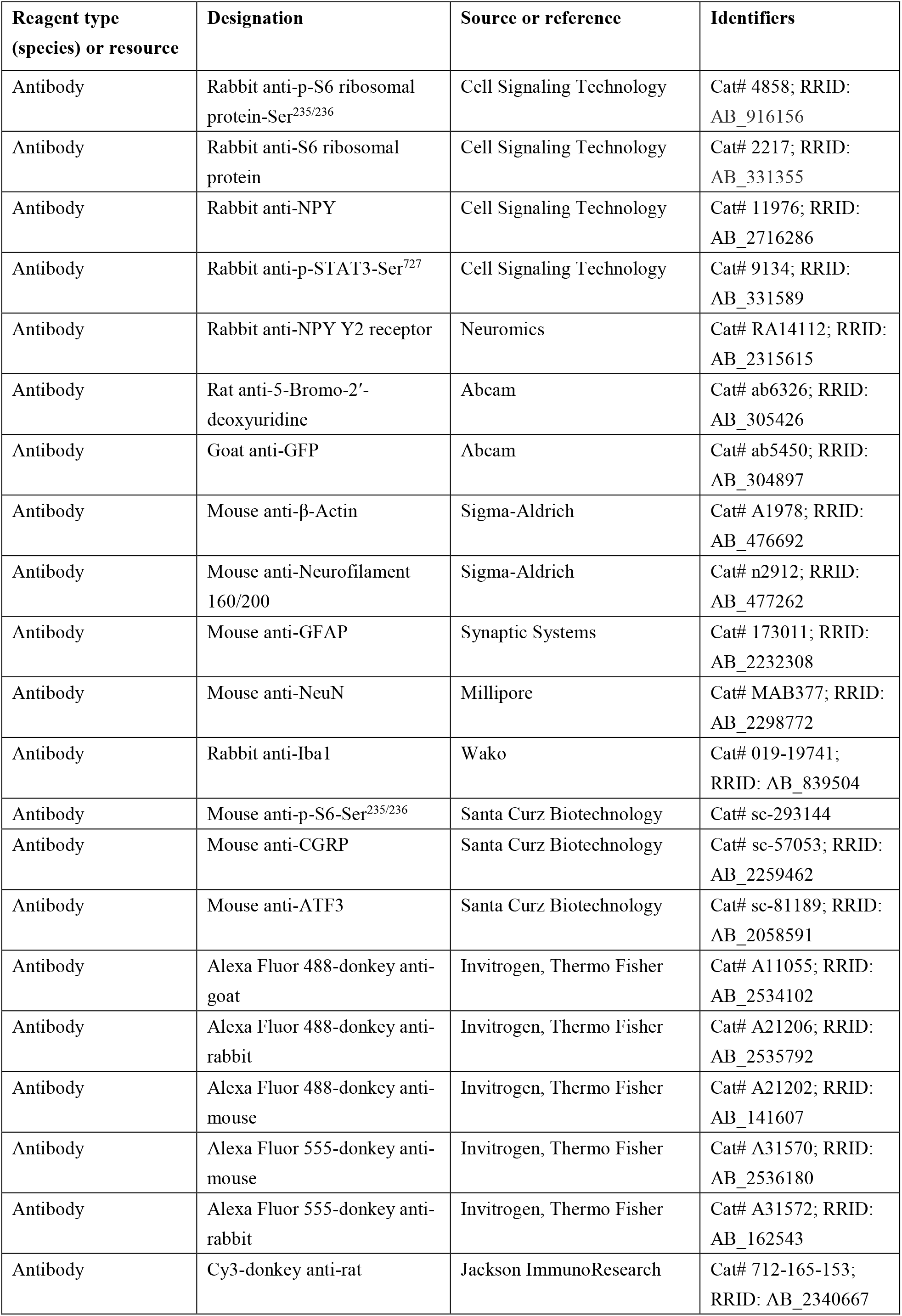
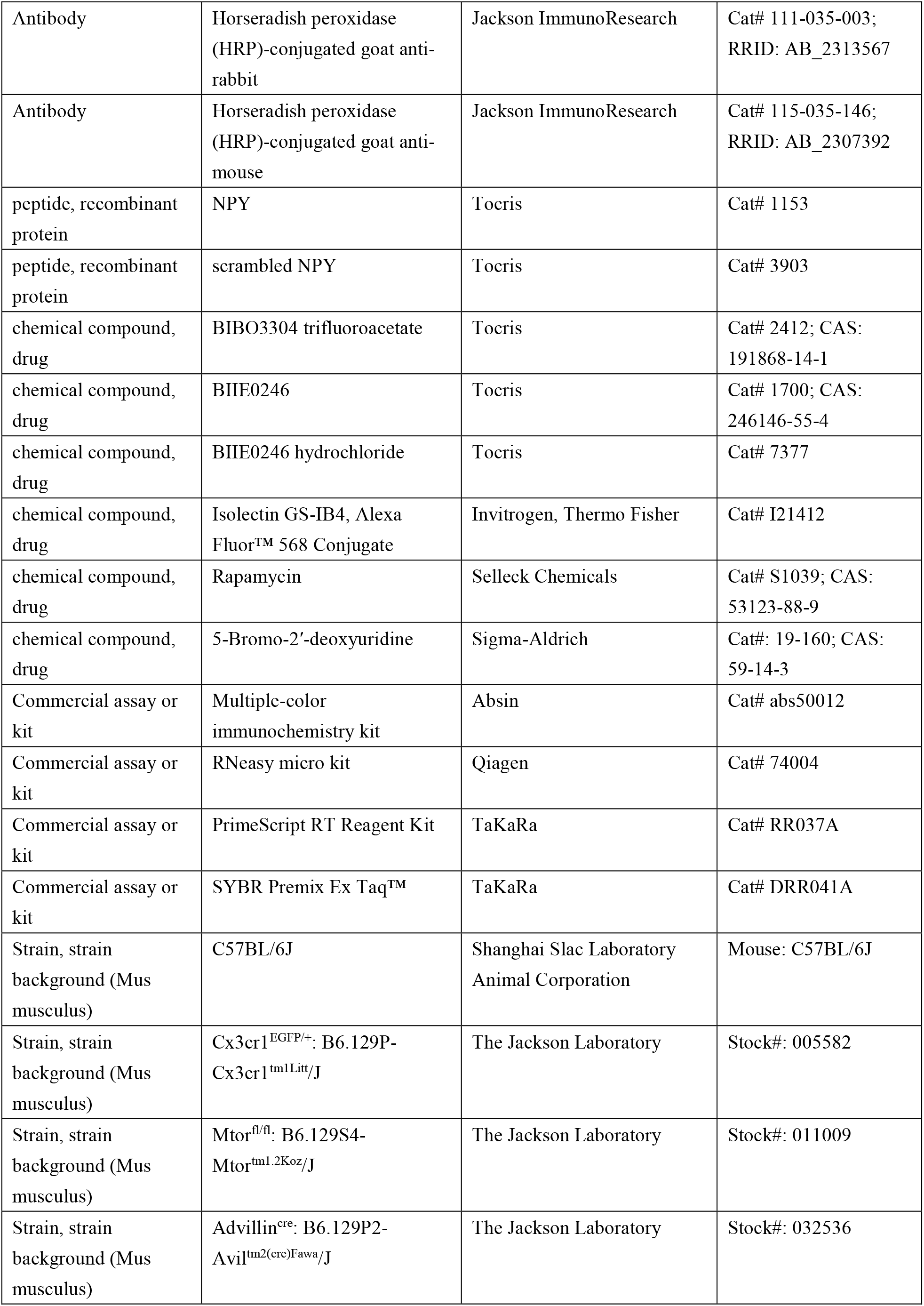
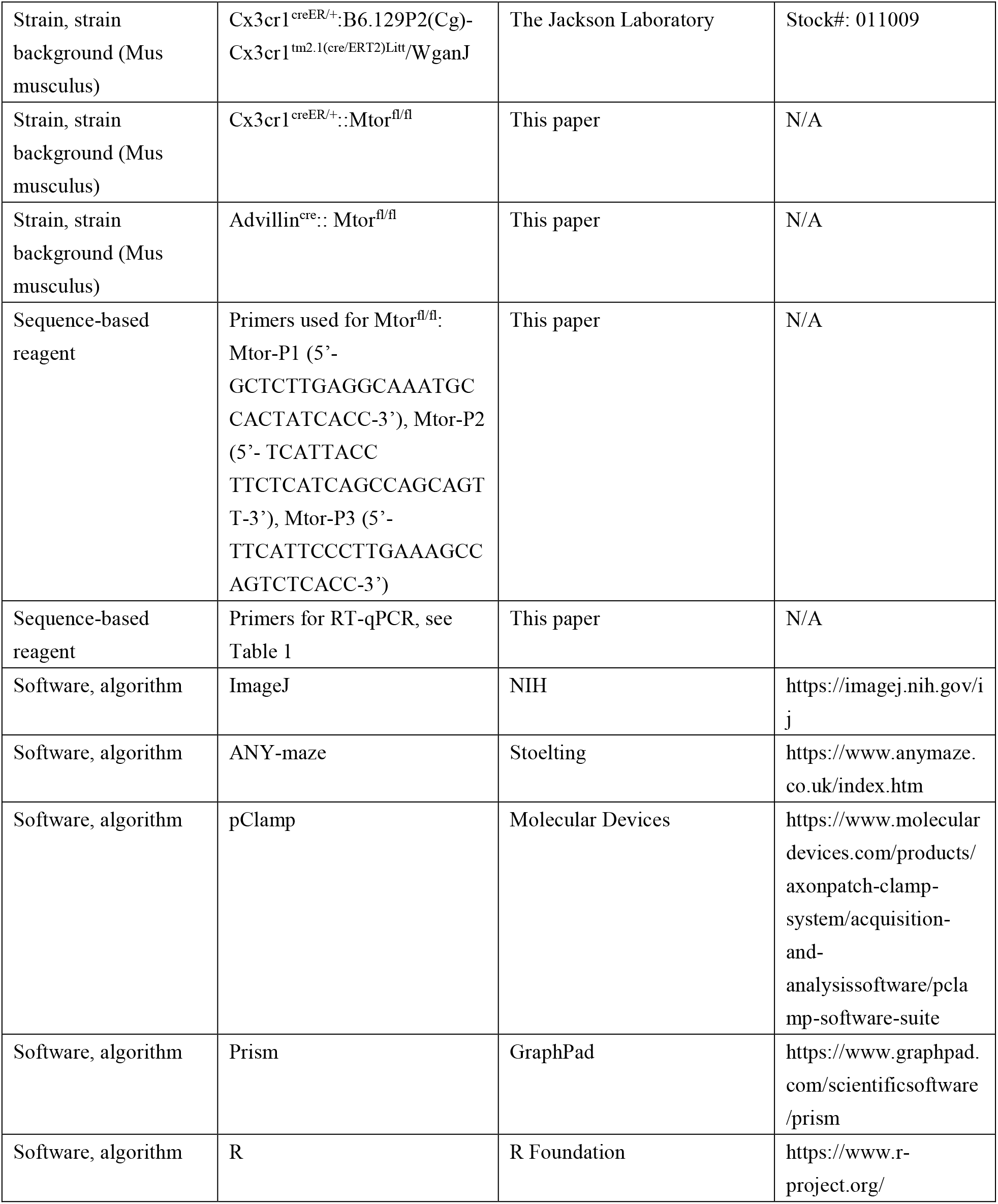

### Whole-cell patch clamp recording

Mice were anesthetized with sodium pentobarbital before sterilized with 75% alcohol. L4 and L5 DRGs were carefully collected on ice and digested with collagenase IV (0.2 mg/mL, Cat# LS004188, Worthington, NJ, USA) and dispase-II (3 mg/mL, Cat# D4693, Sigma-Aldrich) for 60 min at 37 °C. The cell suspension was centrifuged at 500 g for 10 min through a cushion of 15% BSA (Cat# A9205, Sigma-Aldrich) in order to eliminate most of the cellular debris. The cell pellet was resuspended in Neurobasal medium (Cat# 21103049, Thermo Fisher Scientific) with B27 (Cat# 17504-044, Invitrogen, Thermo Fisher Scientific) and NGF supplement (50 ng/mL, Cat# 13257- 019, Gibco, Thermo Fisher Scientific) and seeded onto glass coverslips coated with poly-D-lysine (Cat# P7280, Sigma-Aldrich) and cultured in 5% CO_2_ incubator at 37 °C for at least 2 h before recording. For drug treatment, cultured DRG neurons were incubated with 300 nM NPY for 30 min before recording. To antagonize Y1R or Y2R, BIBO3304 trifluoroacetate (BIBO3304, 1 μM) or BIIE0246 hydrochloride (1 μM) was replenished respectively into medium 30 min before NPY addition.

Whole-cell patch clamp recordings were carried out at room temperature using a Multiclamp 700B amplifier (Molecular Devices, CA, USA). The resistances of borosilicate glass electrodes were measured ranging from 3 to 5 MΩ. The intracellular pipette solution contained (in mM) 135 K-gluconate, 6 NaCl, 10 HEPES, 0.5 EGTA, 10 Na_2_-phosphocreatine, 4 Mg-ATP, 0.3 Na_2_-GTP, and was adjusted to pH 7.2 using KOH. The extracellular solution was composed of (in mM) 150 NaCl, 5 KCl, 2.5 CaCl_2_, 1 MgCl_2_, 10 HEPES, and 10 glucose, and was adjusted to pH 7.4 by NaOH. Action potential firing and resting membrane potential (RMP) were recorded from small-diameter neurons (< 20 μm). Data were collected from neurons with stable RMP negative than -40 mV. Action potentials were evoked by current injection steps. Data were digitized with Digidata 1440A (Molecular Devices), and analyzed by *pClamp* software (Version 10.6, Molecular Devices).

### Behavioral tests

The following behavioral tests were conducted in a blinded manner and during daytime (light cycle). For all experiments, experimenters were blinded to genotypes or experimental manipulation. All the apparatuses and cages were sequentially wiped with 70% ethanol and ddH_2_O then air-dried between stages.

### von Frey tests

*von Frey* tests were used to evaluate 50% paw withdrawal threshold (50% PWT) during the light cycle. In brief, individual mouse was habituated in an opaque plexiglas chamber on a wire mesh platform for 30 minutes prior to test. Testing was performed using a set of von Frey filaments (0.008-2 g, North Coast Medical, CA, USA). Each filament was applied to the lateral part of plantar surface of the mouse hind paw vertically for up to 3 seconds from the bottom. Positive response was determined as a sharp withdrawal, shaking or licking of the limb. The 50% PWT was determined by the up-down method (Dixon, 1965). Test was carried out at 1 day before SNI (baseline) and at 1, 3, 5, and 7 days post-surgery.

### Hargreaves tests

Thermal sensitivity was examined using Hargreaves radiant heat apparatus (IITC Life Science, CA, USA). The basal paw withdrawal latency was adjusted to 9-12 s, with a cutoff of 20 s to avoid tissue damage.

### Hot plate tests

Mice were placed on the hot plate (IITC Life Science) at 50, 52 or 56 °C and the reaction time was scored when the animal began to exhibit signs of pain avoidance such as jumping or paw licking. Animals that did not respond to the noxious heat stimulus after 40 s were removed from the plate.

### Acetone tests

For cold allodynia, 20 μL acetone was applied to the ventral surface of a hind paw, and then the mouse’s response was observed for 60 s. The duration of the mouse responding to acetone, such as withdrawal or flick of the paw, was recorded.

### Rotarod tests

A Rotarod system (Panlab, Spain) was used to assess motor function. Mice were tested in 3 separated trials with a 10 min interval. During the tests, the speed of rotation was accelerated from 4 to 40 rpm over a 5 min period. The falling latency was recorded.

### Open field tests

Mice were placed in the middle of a novel open field arena (45 cm length × 45 cm width × 30 cm height) under normal light conditions. Using *ANY-maze* software (Stoelting, IL, USA), the distance the animal walked in 10 min was recorded.

### Conditioned place aversion (CPA) tests

CPA experiments were conducted in a two-chamber device (50×25 cm) at day 15 post SNI. The CPA protocol included pre-conditioning (baseline), conditioning, and post-conditioning phases (10 min during each phase). Animals spending > 500 s or < 100 s of the total time in either chamber in the pre-conditioning phase were eliminated from further analysis. Immediately following the pre-conditioning phase, the mice underwent conditioning for 10 min. During conditioning, one of the two chambers was paired with the mechanical stimuli. The mechanical stimulus was repeated every 10 s with a 0.16 g von Frey hair on the left hind paw when the mouse enters into the condition chamber. During the post-conditioning phase, the animals did not receive any stimuli and had free access to both compartments for a total of 10 min. Animal movements in each of the chambers were recorded, and the time spent in both chambers was analyzed using *Any-maze* software. Difference scores were defined as post-conditioning time subtracted from preconditioning time spent in the stimuli-paired chamber.

### Data availability

Sequencing data have been deposited in GEO under accession codes GSE184014. All data generated or analysed during this study are included in the manuscript and supporting file; Source Data files have been provided for Figures 1, 2, 3, 4, 6, 7, and 8, and corresponding supplementary figures.

### Statistical analysis

Statistical analyses were performed using *GraphPad Prism* (Version 8.0.1, CA, USA). Quantitative measurements are presented as mean ± standard errors of the means (SEM). Measurements lies outside two standard deviations (SD) are excluded. Statistical differences in comparison to the control group were analyzed using paired or unpaired t-tests as appropriate. One-way (for multiple comparisons) or two-way ANOVA (for multiple time points) with Bonferroni’s *post hoc* tests were used for experiments with more than 2 groups. Significance was considered with *p* value < 0.05. Regarding replication, every mouse represents a replicate, and the number of replicates and additional information on statistics (sample sizes, tests and *p* values) are mentioned for each experiment in the figure legend.

### Study approval

All experiments were conducted in the Zhejiang University School of Medicine. The use and care of animals in all experiments followed the guidelines of The Tab of Animal Experimental Ethical Inspection of the First Affiliated Hospital, College of Medicine, Zhejiang University (No. 2017054).

## Acknowledgements

We thank Dr. Zhenzhong Xu for discussions and suggestions on experimental design and technical support, Drs. Chong Liu and Liang Wang for generously providing transgenic mice, and Drs. Kaiyuan Li, Zhongya Wei, and Xiaobo Wu for the help with whole-cell patch clamp recording. We are grateful to research assistants Sanhua Fang and Daohui Zhang at the Core Facilities of Zhejiang University Institute of Neuroscience.

## Funding

This work was supported by National Natural Science Foundation of China (81772382 and 32070974), National Key Research and Development Program of China (2017YFA0104200), Science Technology Department of Zhejiang Province (2020C03042), the Fundamental Research Funds for the Central Universities of China (2019FZA7009), and the Central Universities granted by Zhejiang University (No. 2021FZZX005-29).

## Competing interests

The authors have declared that no conflict of interest exists.

## Author contributions

Lunhao Chen: Conceptualization; Resources; Data curation; Software; Formal analysis; Investigation; Visualization; Methodology; Writing – original draft; Project administration; Writing - review and editing. Yaling Hu: Conceptualization; Data curation; Software; Formal analysis; Investigation; Methodology; Writing – original draft; Writing - review and editing. Siyuan Wang: Data curation; Investigation; Methodology; Writing - original draft. Kelei Cao: Data curation; Software; Formal analysis. Weihao Mai: Software. Weilin Sha: Data curation. Ma Huan: Writing - review and editing. Yong-Jing Gao: Methodology; Writing - review and editing. Shumin Duan: Resources; Supervision; Funding acquisition; Methodology. Yue Wang: Resources; Data curation; Formal analysis; Supervision; Funding acquisition; Validation; Methodology; Writing - original draft; Project administration; Writing - review and editing. Zhihua Gao: Conceptualization; Data curation; Formal analysis; Supervision; Funding acquisition; Validation; Investigation; Writing - original draft; Project administration; Writing - review and editing.

---

**Figure 1-figure supplement 1.**
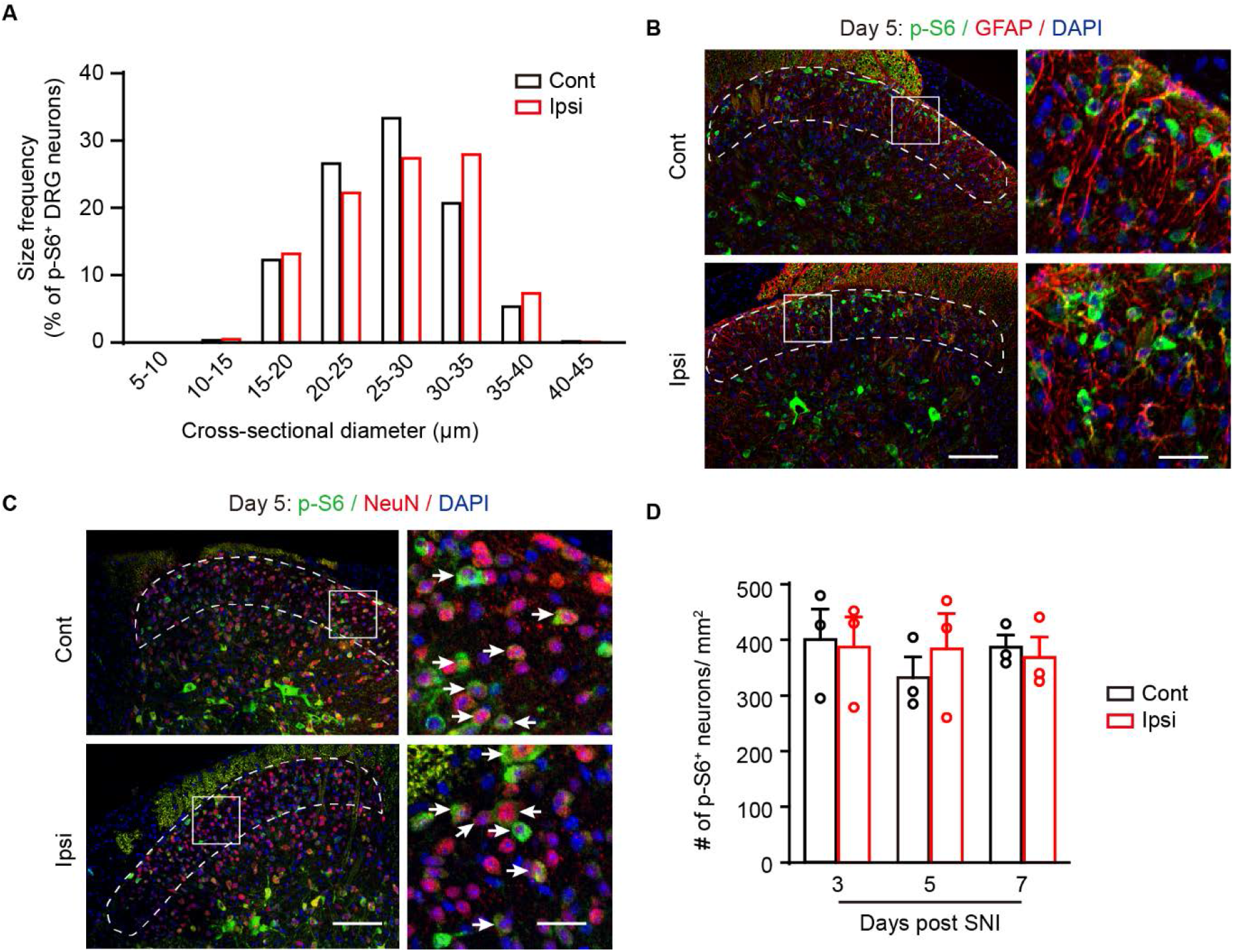
Characterizing of p-S6 staining with different markers in DRG or SDH after SNI. (**A**) Size frequency distribution patterns of p-S6 positive neurons from contralateral and ipsilateral DRG at day 3 post SNI (n=579 and 718 neurons from 3-4 mice respectively). (**B**) Representative images of p-S6 and GFAP in superficial SDH (dotted regions) at day 5 after SNI. Boxes show the region with magnification. (**C**) Representative images of p-S6 and NeuN immunolabeling (arrows) in superficial SDH (dotted regions) at day 5 after SNI. Boxes show the region with magnification. Scale bars, 100 μm and 20 μm for lower- and higher-magnification images, respectively. (**D**) Quantitation of p-S6^+^ neurons in superficial SDH at day 3 to 7 after SNI (n=3 mice per time point). Values are means ± SEM. Two-way ANOVA followed by Bonferroni’s *post hoc* tests among groups. Scale bars, 100 μm and 20 μm for lower- and higher-magnification images, respectively. Ipsi, ipsilateral; Cont, contralateral. **Figure 1-figure supplement 1-source data 1.** Source data used to generate Figure A and D.

**Figure 2-figure supplement 1.**
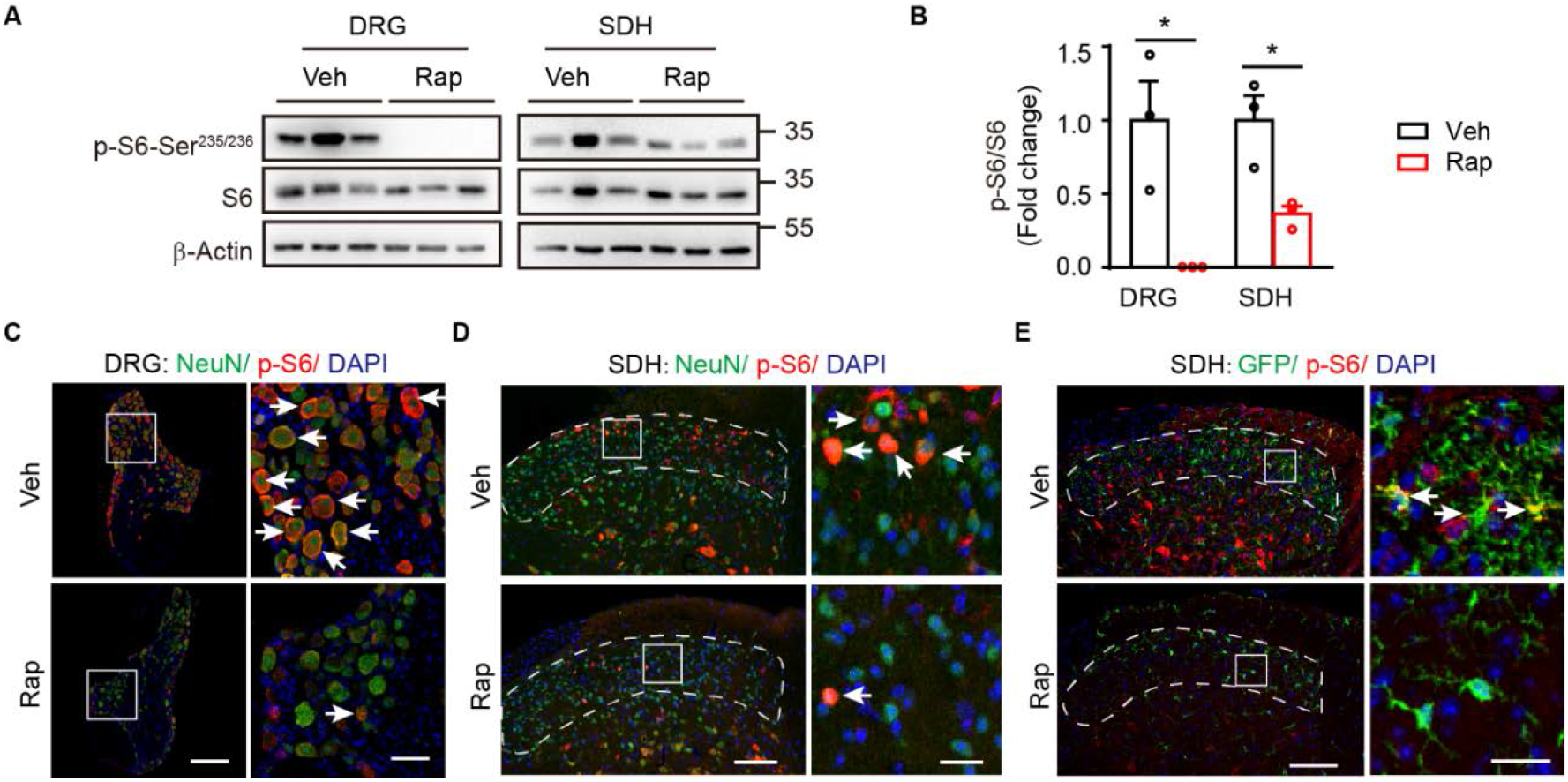
Administration of rapamycin suppresses mTOR activation. (**A**) Representative blots indicating the decreased p-S6 levels in the ipsilateral DRG and SDH following 7-day continuous intraperitoneal (i.p.) injection of rapamycin or vehicle in *Mtor^fl/fl^* mice. (**B**) Quantitation of p-S6/S6 in DRG and SDH following rapamycin treatments (n=3 mice per group). (**C-E**) Co-immunostaining of p-S6 with NeuN or GFP in ipsilateral DRG (**C**) and SDH (**D, E**) after i.p. administration of vehicle or rapamycin (arrows indicating p-S6^+^ cells) at day 7 after SNI. Boxes show regions with magnification. Scale bars, 100 μm and 20 μm in (**C**), and 200 μm and 50 μm in (**D, E**) for lower- and higher-magnification images, respectively. Values are means ± SEM. * *p*<0.05 and ** *p*<0.01 versus Veh, unpaired student t-tests (**B**). Rap, rapamycin; Veh, vehicle; BL, baseline; SDH, spinal dorsal horn. **Figure 2-figure supplement 1-source data 1.** Source data used to generate Figure B. **Figure 2-figure supplement 1-source data 2.** Original pictures of the western blots presented in Figure A.

**Figure 3-figure supplement 1.**
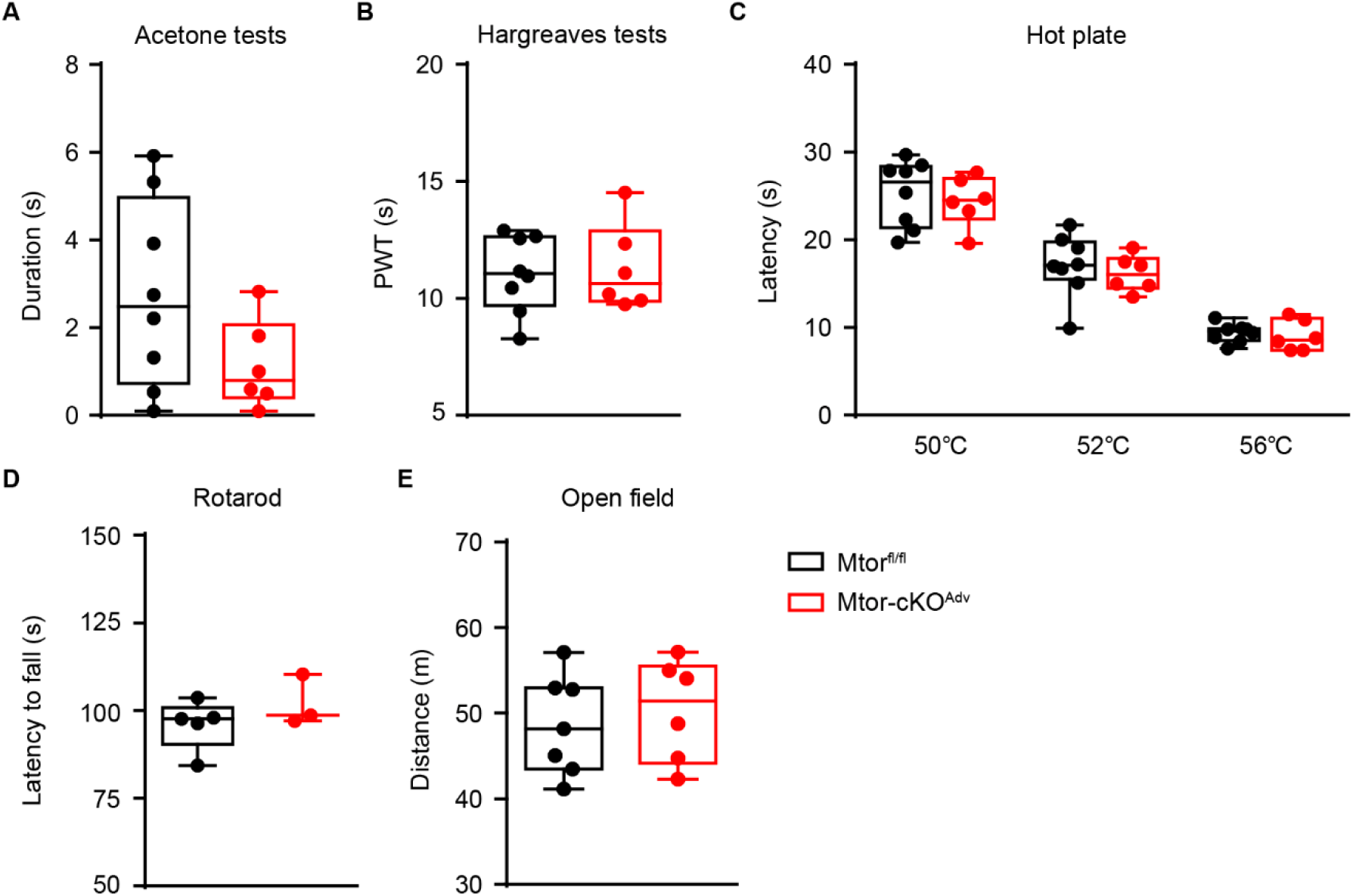
Sensory functions and motor activities are comparable in *Mtor^fl/fl^* and *Mtor-cKO^Adv^* mice at basal state. (**A**) Acetone tests (n=6-8 mice per group). (**B**) Hargreaves tests (n=6-8 mice per group). (**C**) Hot plate tests (n=6-8 mice per group). (**D**) Rotarod tests (n=3-5 mice per group). (**E**) Open field tests in *Mtor^fl/fl^* and *Mtor-cKO^Adv^* mice (n=6-7 mice per group). Values are means ± SEM. Unpaired student t-tests (**A, B, D, E**) and two-way ANOVA followed by Bonferroni’s *post hoc* tests among groups (**C**). **Figure 3-figure supplement 1-source data 1.** Source data used to generate Figure A-E.

**Figure 4-figure supplement 1.**
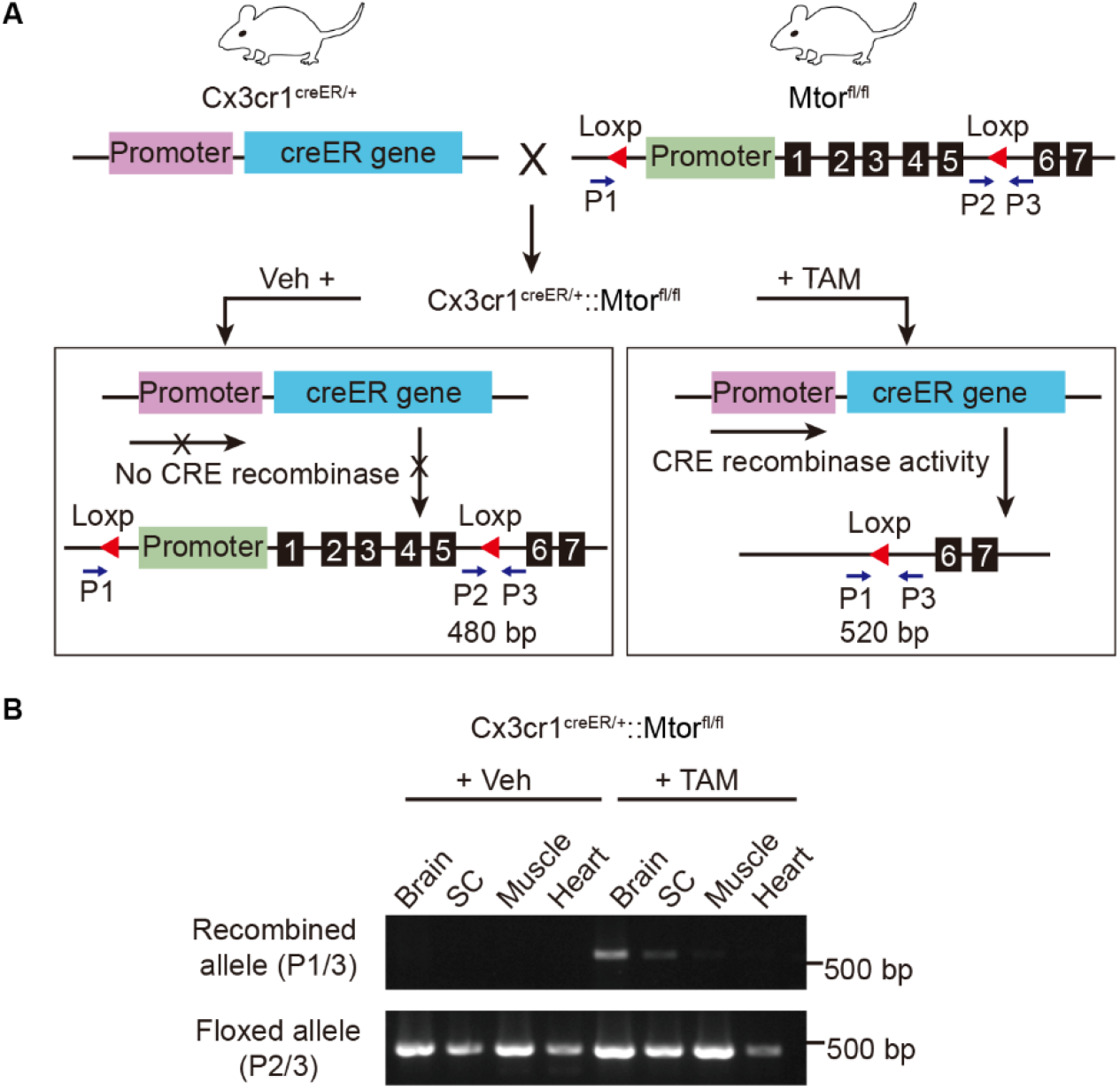
Stratagem for generating *Mtor-cKO^MG^* mice. (**A**) Schematic showing the generation of *Mtor-cKO^MG^* mice. Exons 1-5 of the *Mtor* gene is flanked by *loxP* sites and excised in microglia expressing Cx3cr1-Cre recombinase after TAM administration. The position of P1, P2 and P3 primers and the size of the DNA segments amplified by primer pairs are illustrated. (**B**) Agarose gel electrophoresis of P1, P2 and P3 PCR products showing that Cre-mediated recombination is specifically occurred in the central nervous system (brain and spinal cord), but not in other peripheral tissues (muscle or heart). TAM, tamoxifen; Veh, vehicle. **Figure 4-figure supplement 1-source data 1.** Original pictures of the blots presented in Figure B.

**Figure 5-figure supplement 1.**
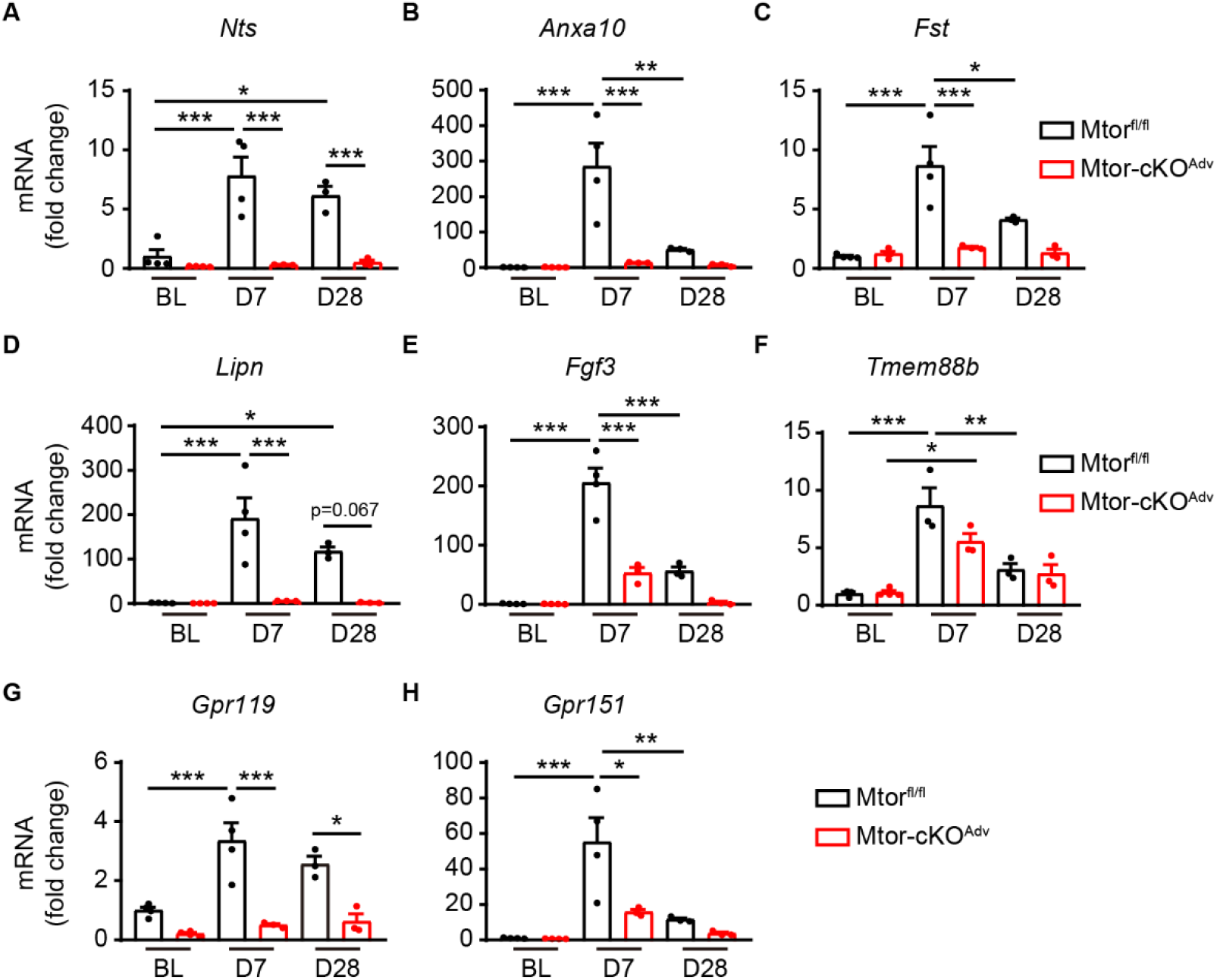
Quantitative RT-PCR of downregulated DEGs identified in RNA sequencing. (**A**) *Nts*, *Neurotensin*; (**B**) *Anxa10*, *Annexin A10*; (**C**) *Fst*, *Follistatin*; (**D**) *Lipn*, *Lipase family member N*; (**E**) *Fgf3*, *Fibroblast growth factor 3*; (**F**) *Tmem88b*, *Transmembrane protein 88b*; (**G**) *Gpr119*, *G protein-coupled receptor 119*; (**H**) *Gpr151*. n=3-4 mice per time point per group. * *p*<0.05, ** *p*<0.01, *** *p*<0.001, one-way ANOVA followed by Bonferroni’s *post hoc* tests among groups. BL, baseline; D, day; DEGs, differentially expressed genes. **Figure 5-figure supplement 1-source data 1.** Source data used to generate Figure A-H.

**Figure 6-figure supplement 1.**
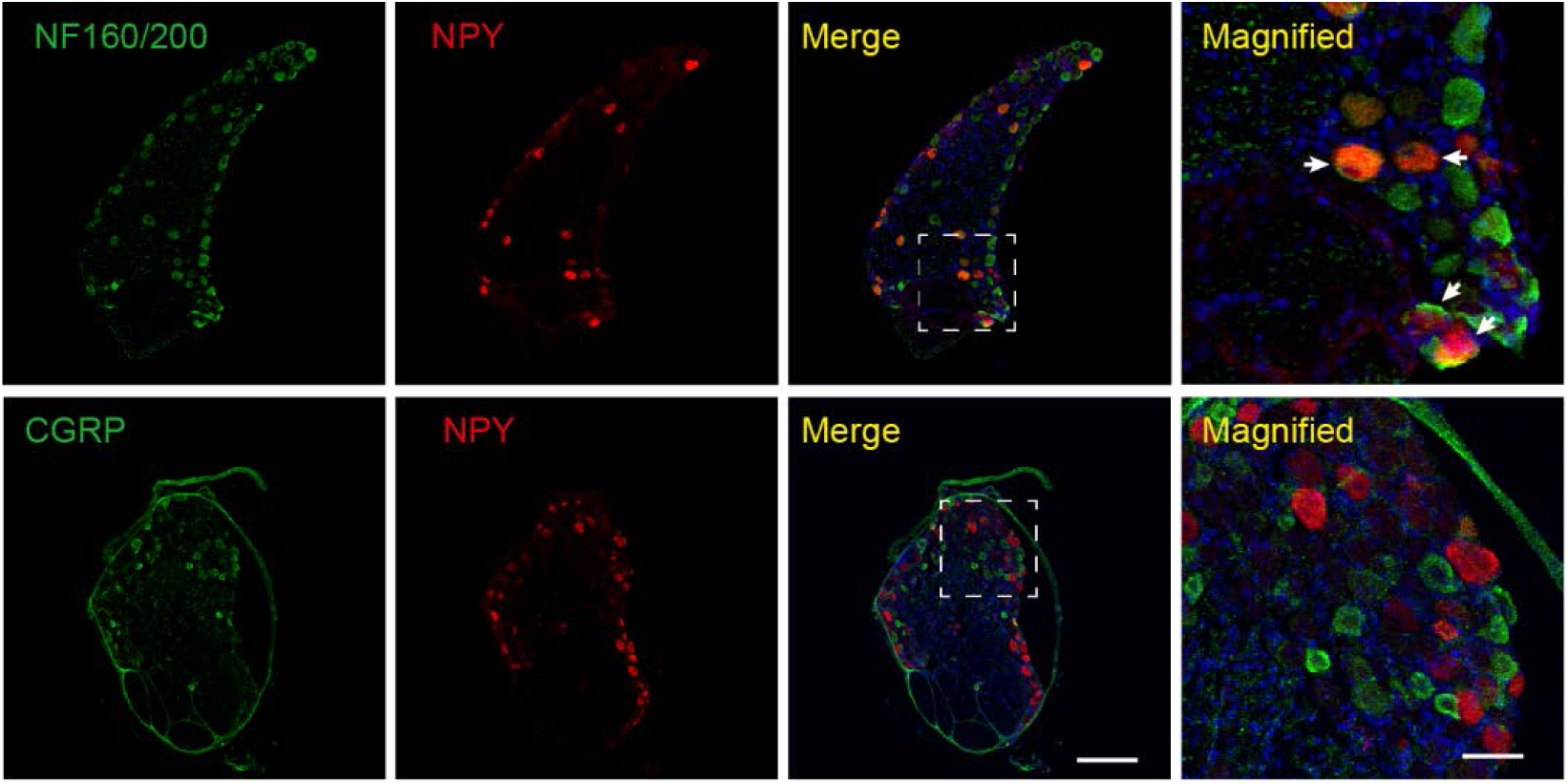
NPY was expressed in large-sized mechanoreceptors. Representative images of NPY and NF160/200 or CGRP staining in the injured DRG. Arrows indicating both NPY^+^ and NF160/200^+^ neurons. Boxes show regions with magnification. Scale bars, 200 μm and 50 μm for lower- and higher-magnification images, respectively.

**Figure 7-figure supplement 1.**
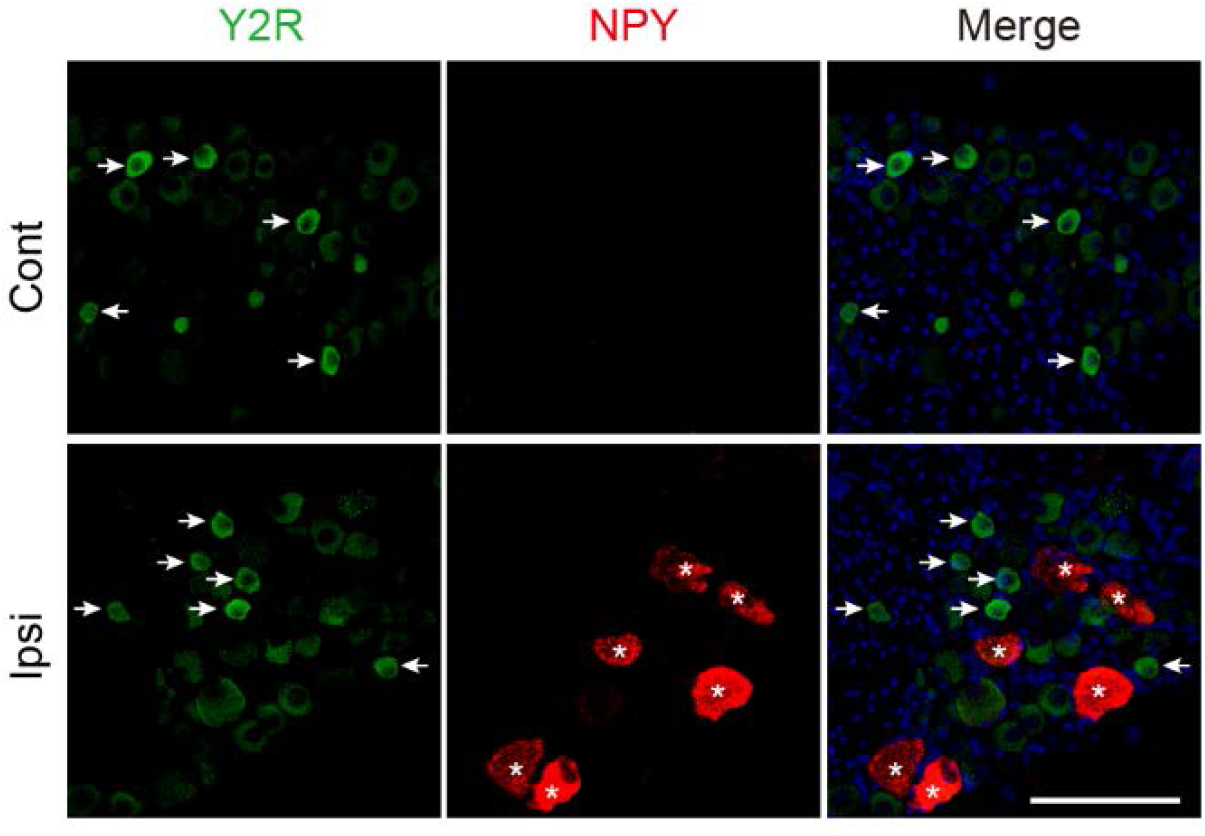
Distinct expression pattern of NPY (*) and Y2R (arrows) by immunofluorescence analysis. Scale bar, 100 μm.

## Notes

### Competing Interest Statement

The authors have declared no competing interest.

